# Between Friends and Foes: Evolutionary Diversification in Mutualistic-Antagonistic Networks

**DOI:** 10.64898/2026.03.16.712075

**Authors:** Felix Jäger, Nicolas Loeuille, Youssef Yacine, Korinna T. Allhoff

## Abstract

Biotic interactions can drive evolutionary diversification, but the underlying mechanisms differ depending on the type of interaction. For instance, Ehrlich and Raven’s escape-and-radiate coevolution provides a pathway of diversification in antagonistic interactions, whereas in mutualistic networks, coevolution is hypothesized to result in trait convergence rather than diversification. The combined effect of mutualism and antagonism on diversification remains unclear, even though organisms naturally engage in multiple types of interactions simultaneously. Using an eco-evolutionary simulation model, we investigate diversification in tripartite ecological networks such as plant-pollinator-herbivore networks. We find that diversification patterns vary according to the way mutualism and antagonism are connected on the trait level. If the two interactions are governed by uncorrelated plant traits, we observe little diversification in the mutualistic and substantial diversification in the antagonistic subnetwork. By contrast, if the same plant trait mediates both mutualism and antagonism (an example of ‘ecological pleiotropy’), diversification rates in all guilds become interdependent. In this case, even the mutualistic guild diversifies considerably when antagonism is strong, while strong mutualism restricts diversification also in the antagonistic guild. Our study underlines that the inclusion of multiple interaction types is necessary to advance our understanding of evolutionary dynamics in ecological networks.

## Introduction

Since Darwin’s ‘On the Origin of Species’, the idea that biotic interactions can drive evolutionary diversification has been prevalent in ecological and evolutionary research. By now, mounting empirical evidence from both palaeological and neontological studies has confirmed associations between ecological interactions and lineage diversification rates, using various approaches ranging from phylogenetic comparative methods to community phylogenetics [1]. Still, the specific mechanisms that drive the observed patterns mostly remain unclear [2, 3].

Different types of biotic interactions are supposed to shape diversification patterns in contrasting ways, via mechanisms that act across multiple evolutionary scales. A large body of work has focused on competition, which, by favouring dissimilar phenotypes, can cause phenotypic diversification within species as well as macroevolutionary adaptive radiation [4]. A classic example are Darwin finches on the Galapagos Islands, where competition-induced character displacement in beak size entailed rapid speciation [5]. Similarly, antagonism is thought to enhance diversification [2, 6]. Triggered by Ehrlich and Raven’s hypothesis on escape-and-radiate coevolution [7], a large body of work focused on the co-diversification of herbivores and plants [8, 9, 10]. For instance, phylogenetic studies revealed that changing levels of investment in defensive traits contributed to macroevolutionary diversification in the plant genera *Asclepias* [11, 12] and *Bursera* [13, 14]. Empirical evidence also exists for antagonistic coevolution driving within-species diversification, e.g., in bacteria-bacteriophage interactions [15, 16], and speciation, e.g., in North-American crossbills [17, 18, 19].

By contrast, implications of mutualism for diversification are less clear [20]. Several micro- and macroevolutionary studies (for instance, using phylogenetic comparative methods) have suggested links between mutualism and diversification, notably pollinator-driven diversification in angiosperms or radiation of mutualistic partners into novel adaptive zones [3, 21, 22, 23, 24, 25]. However, it often remains unclear whether mutualistic coevolution alone or an additional mechanism, such as similarity-based competition was responsible for diversification [6, 10]. Even in obligate pollination/seed-predation mutualisms such as between figs and fig wasps, mutualistic coevolution could not unequivocally be identified as the driver of potential co-speciation [10, 26, 27]. Theoretical work often predicts that mutualistic coevolution prevents diversification by imposing stabilising selection [6, 28, 29, 30, 31]. Stabilising selection exerted by mutualistic partners is also found in empirical studies and can, for instance explain long-term morphological stasis [32].

In nature, organisms are never involved in only one type of biotic interaction [33]. For example, plants engage in mutualistic interactions with their pollinators, but also suffer from herbivory. There exists evidence from theoretical models as well as from empirical work that combining mutualistic and antagonistic interactions can have intricate effects on selection patterns [reviewed in 34]. For example, floral traits such as olfactory or visual cues might attract pollinators and herbivores, creating an ecological trade-off in which benefits of the plant signal due to increased pollination entail costs in terms of herbivory [35, 36, 37, 38]. Resulting conflicting selection pressures can lead to disruptive selection that would not be present if only one type of interaction were considered [39]. However, on a network scale, the joint influence of different types of interactions on evolutionary diversification has hardly been investigated before [but see 40, 41, for a multiple-interaction-type network assembly model without explicit trait dynamics].

We hypothesise that a key determinant of diversification patterns in systems combining antagonism and mutualism is the presence or absence of ‘ecological pleiotropy’ [*sensu* 42]. In analogy to genetic pleiotropy, ecological pleiotropy denotes a situation in which the same trait governs two distinct interactions of an organism. Examples include flower scent or morphology that simultaneously attracts herbivores and pollinators, as mentioned above, or fruit shape that simultaneously determines potential seed dispersers and antagonistic frugivores. A very similar phenomenon occurs if the two traits governing mutualism and antagonism are strongly correlated [43]. Such a correlation can, for instance, be a direct consequence of genetic pleiotropy [43, 44], or arise from an allocation trade-off, such as in plants that can use secondary metabolites either for herbivore defence or the attraction of symbiotic fungi [45]. At the other extreme, the two interactions could be mediated by completely independent traits. For instance, the extent of herbivory may be mainly determined by plant defence compounds that are not correlated with the floral traits relevant for pollination. In this case, no ecological trade-off is to be expected, and the evolutionary dynamics of both interactions are less tightly connected.

In this study, we model the eco-evolutionary assembly of a tripartite ecological network that contains mutualistic and antagonistic interactions. An illustrative example may be plant-pollinator-herbivore networks, but results are also applicable to other types of networks, e.g., plants, seed dispersers and antagonistic frugivores [46, 47, 48] or plants, symbiotic fungi and herbivores [45, 49]. The model is initialised with only one plant, mutualist and antagonist, respectively; during simulations, the network diversifies through coevolution, based on a mutation-selection process. By applying a theoretical model, we are able to test the effect of different ecological processes in isolation. Thereby, we overcome the fundamental problem in empirical research that many processes can create the same macroevolutionary pattern and contribute to a mechanistic understanding of the influence of ecological interactions on diversification [2, 50].

We use the model to investigate (i) how ecological pleiotropy affects evolutionary diversification in tripartite mutualistic-antagonistic networks and (ii) how the effect of ecological pleiotropy on diversification is modulated by the relative strengths of mutualism and antagonism. We hypothesise that ecological pleiotropy can promote diversification by creating an ecological trade-off and subsequent disruptive selection in the plants, followed by corresponding diversification in the other guilds. Based on previous theoretical results, we expect this effect to hinge on a balance of antagonism and mutualism strengths [39].

## Methods

We model the eco-evolutionary emergence of a tripartite ecological network. Rather than structural features of the resulting networks, we here focus on the diversification process itself and how it is influenced by ecological pleiotropy. The network consists of three guilds and includes mutualistic interactions between the first and the second guild as well as antagonistic interactions between the second and the third guild. We assume the central guild to be autotrophs, while the outer guilds depend on the resources provided by the central guild. In the following, we refer to the central guild as plants (P) and to the outer guilds as mutualists (M) and antagonists (A), respectively, encompassing a variety of potential empirical systems. Mutualists and antagonists are thus interacting indirectly via their effect on the plants.

We refer to the members of a guild as ‘phenotypes’. We do not assume them to be necessarily distinct species since our model does not focus on the process of speciation. Instead, the phenotypes could also correspond to different variants within a species. The model is initialised with only one phenotype in each guild. Via the mutation-selection process described below, phenotypes are added to and removed from the system, meaning that all guilds can potentially diversify. The model resembles previous dynamical models of ecological network assembly, e.g., food webs [51, 52], bipartite mutualistic networks [6, 53], or unipartite networks combining different types of interaction [40, 54].

In the following, we first present the equations describing community dynamics, before spelling out how traits determine the interaction strengths in the model. Subsequently, we explain the evolutionary algorithm underlying diversification and describe the performed simulation experiments.

### Community dynamics

We model the density dynamics of the phenotypes in the system via coupled differential equations, similar to previous models combining mutualism and antagonism [40, 54, 55, 56]:

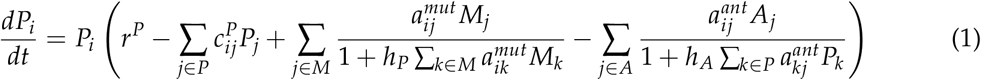

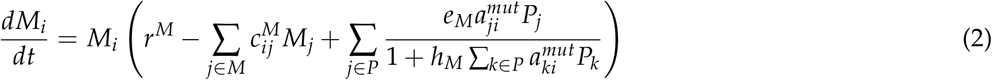

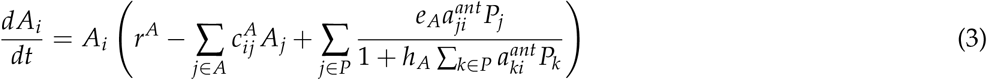

Here, the density of a plant (mutualist, antagonist) phenotype *i* is denoted by *P*_*i*_ (*M*_*i*_, *A*_*i*_). *r*^*P*^, *r*^*M*^ and *r*^*A*^ are intrinsic net growth rates of plants, mutualists and antagonists, respectively. They represent combined birth and mortality rates in the absence of the modelled biotic interactions. Plants are assumed to be autotrophs and can grow without any biotic interactions, meaning *r*_*P*_ *>* 0. In contrast, antagonists and mutualists are heterotrophs and depend on the resources provided by the plants; therefore, *r*^*M*^ and *r*^*H*^ are negative. By assuming that the mutualists and antagonists have no alternative resources to consume, we ensure that all biotic interactions that contribute to the coevolutionary process are made explicit in the model.

The parameters 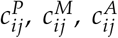 are competition coefficients for intraguild competition between phenotypes *i* and *j*, including intraphenotypic competition (*i* = *j*), and are calculated as described in the section “Intraguild competition” below. The effects of mutualism and antagonism on population dynamics follow a multi-species Holling type II functional response. Parameter 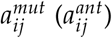 reflects the strength of the mutualistic (antagonistic) interaction between plant phenotype *i* and mutualist (antagonist) phenotype *j*. It corresponds to the attack rate from classical predator-prey models and depends on trait matching as described in the next section. Handling times are *h*_*P*_, *h*_*M*_ and *h*_*A*_ for plants, mutualists and antagonists, respectively. For simplicity, we assume them to be equal and independent of the phenotypes. This is unlikely to hold in specific empirical systems, but the generality of our model does not allow for a finer parametrisation of handling times and the effect of traits on the interguild interactions is already incorporated in the interaction strengths 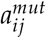 and 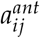. *e*_*M*_ and *e*_*A*_ denote conversion efficiencies of mutualists and antagonists, respectively, and are similarly assumed to be independent of phenotypes.

A Holling type II functional response is frequently used in dynamical models of mutualism to prevent unbounded growth of densities, which can occur with a linear functional response for certain parameter combinations [30, 57, 58, 59, 60]. Note that the functional response for antagonism in equations 1 and 3 always saturates with the plants’ densities as in classical predator-prey models [61], since the antagonist’s benefit is assumed proportional to the plant’s loss. Instead, mutualism most often consists of a reciprocal exchange of services where we assume the mutualist’s benefit to saturate with the number of plants present (see equation 2), while the plant’s benefit saturates with the number of potential mutualist partners (see equation 1) [40, 54, 55, 56].

### Trait matching and ecological pleiotropy

In the model, the processes of interguild mutualism/antagonism as well as intraguild competition are governed by traits of the involved phenotypes. Each mutualist and antagonist phenotype *i* is characterised by a single quantitative trait 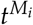 or 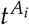, respectively. Plant phenotypes possess two traits 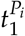 and 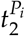 to compare scenarios of ecological pleiotropy with scenarios of uncorrelated interaction traits. These abstract high-level traits aggregate all traits of the respective phenotype that are relevant to mutualism and/or antagonism. In the plant-pollinator-herbivore example, this could include mouth part morphology, scent preferences, volatile cues or phenological traits.

The strength of the interaction between a plant and a mutualist or antagonist increases with similarity of their traits. This trait matching assumption is common in theoretical studies of ecological interactions [e.g., 51, 52, 53, 62, 63, 64, 65] and has empirical foundations in many systems [66, 67, 68]. For example, the matching of proboscis length and corolla length has been repeatedly shown to mediate plant-insect interactions [69, 70] as is the case for bird beak size and fruit size in frugivory [71]. Similarly, matching life-history strategies have been suggested to drive associations of plants and arbuscular mycorrhizal fungi [72]. We use a Gaussian function to mathematically capture trait matching between a plant *i* and a mutualist/antagonist *j*, as depicted in Figure 1A and 1B:

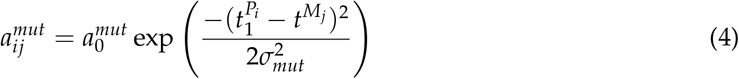

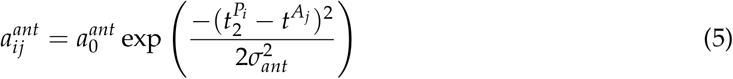

**Figure 1:**
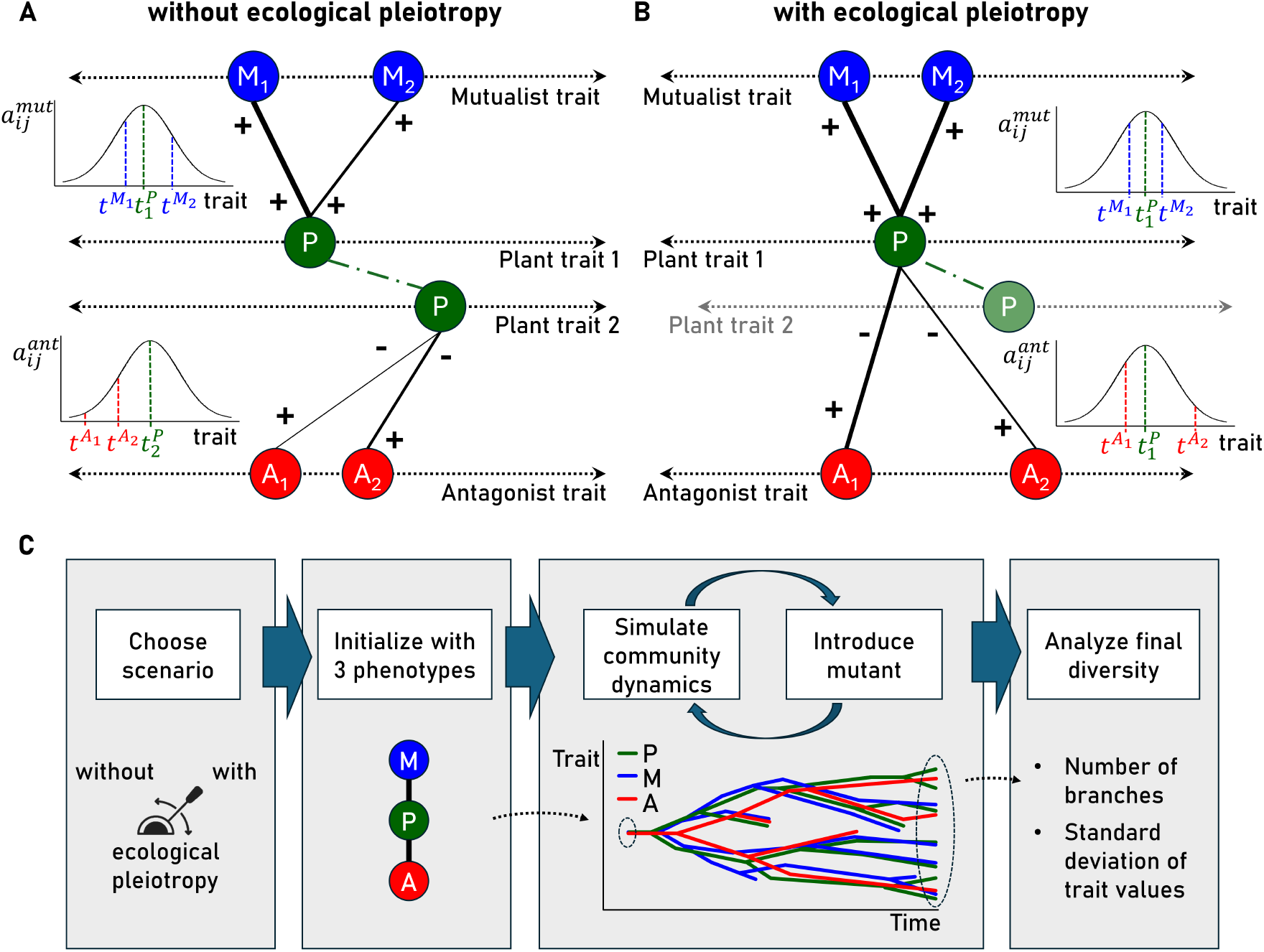
Model overview. **A** Without ecological pleiotropy, the first plant trait determines the strength of mutualism 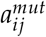, the second plant trait the strength of antagonism 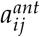. Interaction strengths are maximised when the respective traits are identical. **B** With ecological pleiotropy, the calculation of both interaction strengths is based on the first plant trait. **C** Starting with one phenotype in each guild, the evolutionary algorithm proceeds by alternately simulating the community dynamics and introducing new mutant phenotypes. Final diversity is quantified after 5 billion time steps. All simulations are run with and without ecological pleiotropy.

The parameters 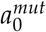 and 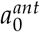 represent the maximal mutualistic/antagonistic interaction strength, the interaction kernel widths *σ*_*mut*_ and *σ*_*ant*_ encapsulate how fast interactions become weak when partner traits differ.

Equations 4 and 5 represent the model version without ecological pleiotropy. The first plant trait modulates the mutualistic, the second plant trait the antagonistic interaction (Fig. 1A). With ecological pleiotropy, 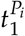 is used instead of 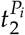 in equation 5. This corresponds to a scenario where one plant trait is decisive for both types of interaction, e.g., when the same floral volatile cues attract both pollinators and herbivores (Fig. 1B) [42]. In this case, the second plant trait is irrelevant for mutualism and antagonism.

### Intraguild competition

Phenotypes compete with other phenotypes of the same guild. We assume competition to be slightly stronger for phenotypes with similar trait values. This is based on limiting similarity theory, assuming that similarity in traits is generally associated with similarity in niches (e.g., temporal or spatial niches) [73, 74, 75, 76, 77, 78]. More straightforward, it can, for instance, reflect competition among pollinators or herbivores for floral resources or interference competition that increases with overlap in visited plants. Our assumption that the same traits that govern interguild interactions also affect intraguild competition (potentially via a trait-dependent carrying capacity) is an ecological pleiotropy in itself and shared by many comparable models in the literature [30, 31, 51, 52, 53, 56, 63, 79]. Nonetheless, we assume that another, larger, proportion of a phenotype’s niche is not explained by these traits. Therefore, our model combines a similarity-based component and a trait-independent component of intraguild competition. The relative contribution of similarity-based competition to total competition for plants (mutualists, antagonists) is described by a parameter *α*^*P*^ (*α*^*M*^, *α*^*A*^), ranging from 0 (no similarity-dependence) to 1 (full similarity-dependence). The baseline value is 0.2, meaning that 80% of the intraguild competition is independent of the interaction traits. The competition coefficients 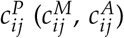 for plant (mutualist, antagonist) phenotypes *i* and *j* are given by the following equations:

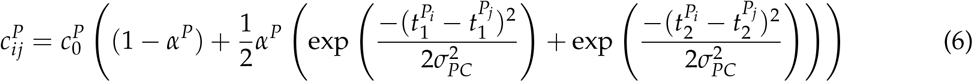

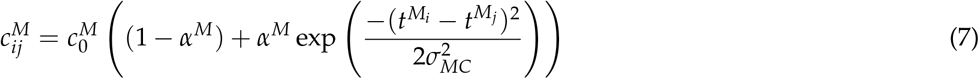

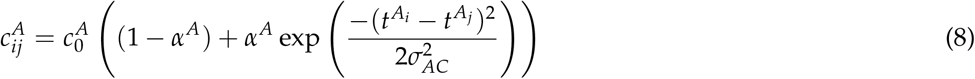

Here, 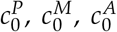 are the maximal competition strengths for plants, mutualists and antagonists, respectively. The widths of the competition kernels *σ*_*PC*_, *σ*_*MC*_ and *σ*_*AC*_ describe how fast competition gets weaker for dissimilar phenotypes. For simplicity, we assume the competition kernel widths as well as the proportions of similarity-based competition (*α*^*P*^, *α*^*M*^, *α*^*A*^) equal among guilds since we focus on the effects of mutualism, antagonism and ecological pleiotropy rather than competition. For plants, similarity-based competition relies on both traits, even in the case of ecological pleiotropy. The competition terms for each trait are added. This reflects the assumption that the competition arises from different mechanisms [the case of ‘non-substitutable resources’ in 80], i.e., similarity in one trait still evokes competition even if the difference in the other trait is high.

Prior model analysis showed that a basic level of similarity-based competition is necessary for any diversification to happen (see Supplementary Material S1). Beyond that, our results are qualitatively robust to varying the proportion of similarity-based competition (see Supplementary Material S2). This identifies similarity-based competition as a prerequisite, but not the main driver of the patterns we find.

### Evolutionary dynamics

Evolutionary dynamics are incorporated into the model by simulating a mutation-selection process, as explained in the following (see Fig. 1C). The algorithm is inspired by the framework of Adaptive Dynamics [81, 82, 83], according to which mutations are assumed to be small and rare.

The model is initialised with only one phenotype in each guild. Then, equations 1, 2, 3 for the community dynamics are solved numerically. After 1 000 time steps, the first mutation occurs. A parent phenotype is selected at random among all guilds, with probability being proportional to the product of its density and the respective mutation rate *µ*_*P*_ (plants), *µ*_*M*_ (mutualists) or *µ*_*A*_ (antagonists). A new mutant is then created belonging to the same guild as the parent, with very low initial density *N*_0_, subtracted from the parent’s density. Its trait value is drawn from a normal distribution around the parent’s trait value with fixed standard deviation Σ, reflecting the amplitude of mutations. For plant mutants, this is done for both traits at the same time. Subsequently, the time until the next mutation is calculated. We assume mutations to follow a Poisson process, with *µ*_*P*_ (*µ*_*M*_, *µ*_*A*_) being the fixed mutation rates of a plant (mutualist, antagonist) individual. The waiting time until the next mutation is therefore drawn from an exponential distribution with parameter *µ*_*P*_ *N*_*P*_ + *µ*_*M*_ *N*_*M*_ + *µ*_*A*_ *N*_*A*_, where *N*_*P*_, *N*_*M*_ and *N*_*A*_ denote total densities of plants, mutualists and antagonists at the current mutation time. Finally, community dynamics are simulated again until the next mutation, and the whole process is repeated until a maximum number of simulated time steps is reached. Phenotypes whose densities fall below *N*_0_ are considered extinct and removed from the system (a check is carried out at each mutation time, not between mutations). This leaves three possibilities after a new mutant is introduced to the system: it either goes extinct (cannot invade), replaces the resident, or coexists with the resident. The latter case results in diversification of the network.

### Simulation design

All simulation runs were initialised with all traits being equal to zero (corresponding to maximum strength of antagonism and mutualism) and initial densities equal to *N*_0_. The default parameter values are listed in Table 1. In the baseline scenario, for comparability of the different interaction types, all parameters of antagonists and mutualists were chosen to be equal. The simulations were run for 5 *·* 10^9^ time steps. To avoid excessive run times, simulations were stopped prematurely if one of the guilds reached a maximum number of 300 phenotypes.

**Table 1.** Model variables and parameters.

| Symbol | Meaning | Default value | Dimension |
| --- | --- | --- | --- |
| *Variables* |  |  |  |
| $P_i, M_i, A_i$ | plant/mutualist/antagonist phenotype densities | - | individuals · area<sup>-1</sup> |
| $t_1^{P_i}, t_2^{P_i}$ | first/second plant trait | - | trait dimension* |
| $t^{M_i}, t^{A_i}$ | mutualist/antagonist trait | - | trait dimension* |
| *Ecological parameters* |  |  |  |
| $r^P$ | plant intrinsic net growth rate | 10 | time<sup>-1</sup> |
| $r^M, r^A$ | mutualist/antagonist intrinsic net growth rate | -0.01 | time<sup>-1</sup> |
| $c_0^P$ | maximum plant competition coefficient | 10 | area · (individuals · time)<sup>-1</sup> |
| $c_0^M, c_0^A$ | maximum mutualist/antagonist competition coefficient | 1 | area · (individuals · time)<sup>-1</sup> |
| $\alpha^P, \alpha^M, \alpha^A$ | plant/mutualist/antagonist share of similarity-based competition | 0.2 | none |
| $\sigma_{PC}, \sigma_{MC}, \sigma_{AC}$ | plant/mutualist/antagonist competition kernel width | 1 | trait dimension* |
| $e_M, e_A$ | mutualist/antagonist conversion efficiency | 3 | none |
| $h_P, h_M, h_A$ | plant/mutualist/antagonist handling time | 0.1 | time |
| $a_0^{mut}, a_0^{ant}$ | maximum strength of mutualism/antagonism | 1 | area · (individuals · time)<sup>-1</sup> |
| $\sigma_{mut}, \sigma_{ant}$ | mutualism/antagonism kernel width | 2.5 | trait dimension* |
| $N_0$ | extinction threshold/initial mutant density | 10<sup>-5</sup> | individuals · area<sup>-1</sup> |
| *Evolutionary parameters* |  |  |  |
| $\Sigma$ | Mutation amplitude (standard deviation) | 0.05 | trait dimension* |
| $\mu_P, \mu_M, \mu_A$ | Mutation rate of a plant/mutualist/antagonist individual | 2 · 10<sup>-6</sup> | area · (individuals · time)<sup>-1</sup> |

\*The dimension of the traits depends on the study system. For instance, traits related to flower or mouthpart morphology are often measured as length or area, the emission rate of floral volatile organic compounds as mass per mass per time [87].

Diversity was quantified using two different measures that encompass different dimensions of diversity. First, as a measure of phenotype richness, the number of ‘branches’ in each guild was calculated. Branches are collections of phenotypes with close trait values that can be considered functionally equivalent. In theory, in the long run, all but one phenotype in a branch would go extinct due to competitive exclusion. However, in the simulations (and in nature) new mutations often accumulate faster than it takes for slightly weaker competitors to go extinct. Branches were identified via a clustering analysis using the DBSCAN algorithm, which is a widely used density-based non-parametric clustering algorithm [84]. For plants, clusters were calculated in the two-dimensional trait space. Second, as a measure of functional diversity, we computed the weighted standard deviation in traits within each guild [square root of weighted trait variance, 85]. Phenotypes were weighted with their density; for plants, weighted trait variances for both traits were added.

We analysed individual simulation runs and tracked all traits, the number of branches and the total density of each guild through time. Moreover, as a proxy for overall diversification, we recorded both the number of branches and the weighted standard deviation of traits at the end of each simulation run. We then computed the mean and the coefficient of variation of these measures over 20 replicates with different random seeds to account for variation due to stochasticity in mutations.

To analyse the contribution of individual interaction types to diversification patterns, we first run the model for five different scenarios: (i) without mutualism and antagonism, (ii) with antagonism but without mutualism (bipartite antagonistic), (iii) with mutualism but without antagonism (bipartite mutualistic), (iv) with mutualism and antagonism but without ecological pleiotropy and (v) with mutualism and antagonism and with ecological pleiotropy. Here, interactions were excluded by setting the maximum strength of antagonism/mutualism 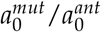 to zero. Subsequently, to examine the influence of relative interaction strengths, we performed simulations for varying combinations of 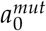 and 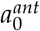, again with and without ecological pleiotropy. Further robustness checks were carried out varying the proportions of similarity-based competition *α*^*P*^, *α*^*M*^ and *α*^*A*^, the net intrinsic growth rates of mutualists and antagonists *r*^*M*^ and *r*^*A*^, the ratio of the mutation rates *µ*_*P*_, *µ*_*M*_ and *µ*_*M*_, and the mutualism/antagonism kernel widths *σ*_*mut*_ and *σ*_*ant*_ (see Supplementary Material S2). In an additional robustness check we examined a model version that relaxes the assumption of neutrality in the absence of interactions by introducing an environmental optimum for the trait values (see Supplementary Material S3).

The model was implemented in the programming language C. The differential equations were numerically integrated using the solver CVODE v7.3.0 from the SUNDIALS library [86].

## Results

### Ecological pleiotropy aligns mutualist and antagonist diversification rates

We start by analysing typical individual simulation runs that illustrate general results on the evolutionary diversification in tripartite networks (see Fig. 2).

**Figure 2:**
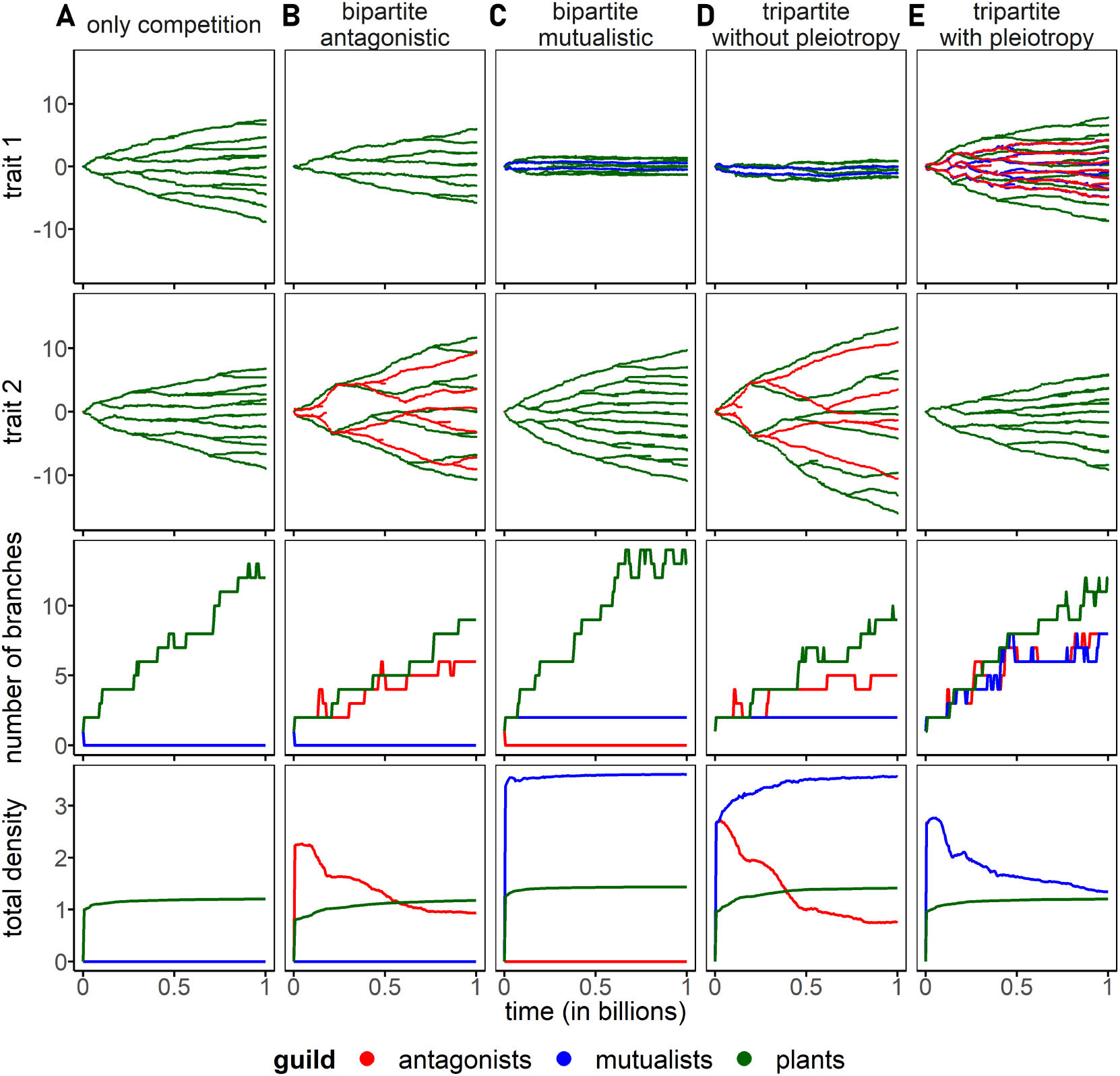
Comparison of typical simulation runs. Columns correspond to scenarios with (A) only competition, (B) competition and antagonism, (C) competition and mutualism, (D) competition, antagonism and mutualism but without ecological pleiotropy, (E) competition, antagonism and mutualism and ecological pleiotropy. Rows show (i) first plant trait, (ii) second plant trait, (iii) the number of branches, and (iv) the total density of each guild over time. The mutualist and antagonist traits are always plotted together with the corresponding plant trait. Parameter values are chosen according to Table 1. For visibility of the trajectories in the trait space, only the first billion time steps are shown.

Without mutualistic and antagonistic interactions (Fig. 2A), both antagonists and mutualists cannot survive since their intrinsic growth rate is negative. Plants, however, diversify in both their traits due to similarity-based competition. Starting from the initial trait value 0, gradually, larger and larger parts of the trait space are filling up with plant branches. The distribution of plant branches exhibits an approximately regular spacing, with an average distance between adjacent branches of 1 to 2, comparable to the width of the competition kernel *σ*_*PC*_ = 1. The total density of plants quickly saturates due to trait-independent competition, which imposes a carrying capacity on each guild as a whole (see bottom row in the figure).

In the bipartite antagonistic network (Fig. 2B), plants and antagonists diversify simultaneously. During the diversification process, plant branches tend to escape the antagonist branches in their second trait, while antagonist branches follow the closest plant branch, similar to classical coevolutionary arms races. Compared to the scenario with only plants, this leads to a faster filling of the outer parts of the trait space and greater distances between the plant branches. The total density of antagonists decreases over time, while plant density saturates at a similar value as in the scenario with only plants. This reflects that several of the plant branches (especially the outer plants) manage to escape their antagonists. Accordingly, the outer plant branches reach higher densities than the central ones (not shown). In our model, higher densities translate into more frequent mutations and thus faster evolution (see Methods). Therefore, the higher densities of the outer branches can, in turn, accelerate their escape from the antagonists in a positive feedback loop. In the first trait, plants diversify less than in the scenario without antagonism. This can be attributed to slightly lower densities in the beginning (reducing the speed of evolution as well as the intensity of similarity-based competition) or to the strong directional selection in the second trait, superimposing selection pressures arising from similarity-based competition in the first plant trait.

In the mutualistic bipartite network (Fig. 2C), the mutualist and first plant trait stay close to the initial trait values and undergo only limited diversification (only two mutualist branches), owing to stabilising selection. However, plants do diversify in their second trait. Due to the ecological benefits from mutualism, plant densities are slightly higher than in the case with only competition. Since higher density translates into higher speed of evolution (see above) as well as more intense competition, this can explain the slight acceleration of plant diversification. Mutualist densities saturate quickly at a high level.

The diversification patterns in the tripartite network without ecological pleiotropy (Fig. 2D) are a combination of the patterns observed in the two bipartite networks. Mutualist and first plant trait do not diversify much, antagonists and second plant trait undergo simultaneous diversification with pronounced escape-and-chase dynamics. Antagonist diversification is slightly slower, and distances between plant branches in the second trait are slightly higher than in the bipartite antagonistic case. This can be explained by the fact that plants can escape faster from the antagonists since their densities are augmented by mutualism.

If ecological pleiotropy is present, meaning that both antagonism and mutualism are governed by the first plant trait, the system shows a qualitatively different pattern (Fig. 2E). All guilds diversify, with the antagonist and mutualist branches following the plant branches, exhibiting very similar trajectories due to the symmetric parametrisation. Plants diversify as in the scenario with competition only, they do not escape from antagonists since mutualists close to the antagonists eliminate the runaway selection. Accordingly, plant branches stay more concentrated in the centre compared to the antagonistic subnetwork in the case without ecological pleiotropy. Also, antagonists and mutualists accumulate in the centre of the trait space, attracted by the high concentration of plant branches there. Hence, more and more outer plant branches remain unoccupied, leading to a simultaneous decline in the total density of antagonists and mutualists (overlapping red and blue lines in the bottom row of the figure).

### Relative interaction strength determines diversification patterns

In a second step, we investigate how the effect of ecological pleiotropy on diversification changes with the relative strength of mutualism vs. antagonism. To this end, the maximum interaction strengths 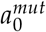 and 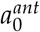 were systematically varied.

With ecological pleiotropy, the relative strength of mutualism and antagonism strongly influences trajectories in the trait space in individual simulation runs (Figure 3). If antagonism strength exceeds mutualism strength, high levels of diversification are observed in all three guilds (3 plots shaded in red in the top left of Figure 3). If instead mutualism strength is higher than antagonism strength (3 plots shaded in blue in the bottom right), diversification in mutualists, antagonists and the first plant trait remains limited. After a brief period of diversification at the beginning, the traits stay approximately constant, close to the initial values. Plants can still diversify in their second trait (not shown).

**Figure 3:**
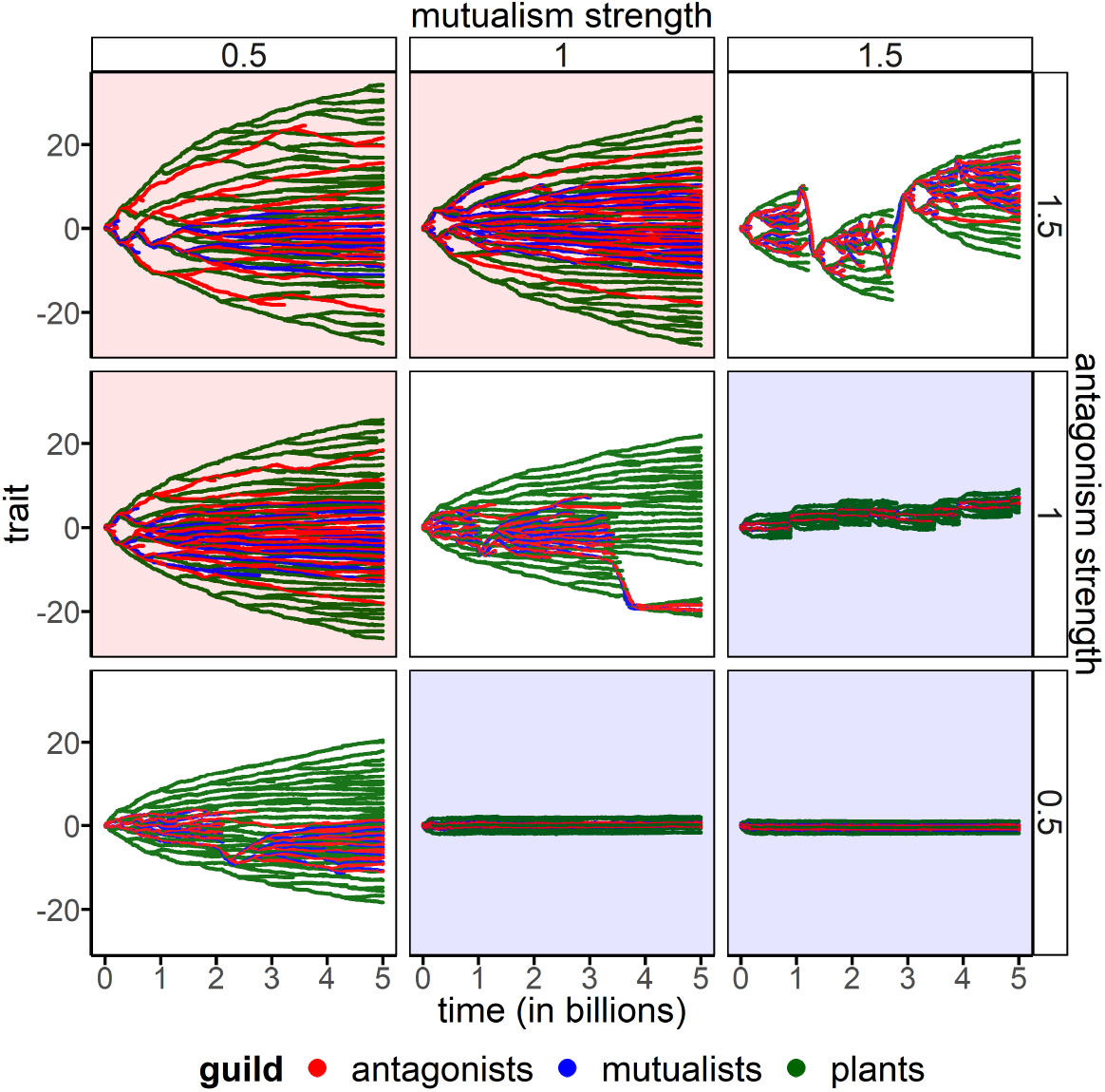
Relative interaction strengths drive evolutionary trajectories with ecological pleiotropy. Time series show antagonist, mutualist and first plant trait for varying strength of antagonism 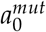 and strength of mutualism 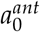 in simulations with ecological pleiotropy. All other parameters are chosen according to Table 1.

Interestingly, if antagonism and mutualism strength match (3 plots without shading on the diagonal in Fig. 3), temporary diversification in all guilds can be observed, interrupted by extinction avalanches. By this term, we denote the extinction of a major portion of branches within a very short time interval. After an extinction avalanche, the diversification process starts afresh. Extinction avalanches can affect all three guilds, as in the top right corner of Figure 3 after around one billion time steps, or a subset of guilds only, as in the bottom left corner after approximately two billion time steps. We provide a possible mechanistic explanation for the occurrence of extinction avalanches in the Supplementary Material S4.

For a systematic evaluation of the effect of ecological pleiotropy on phenotype richness, we analyse the number of branches at the end of a simulation run as a proxy for overall diversification rate (Figure 4).

**Figure 4:**
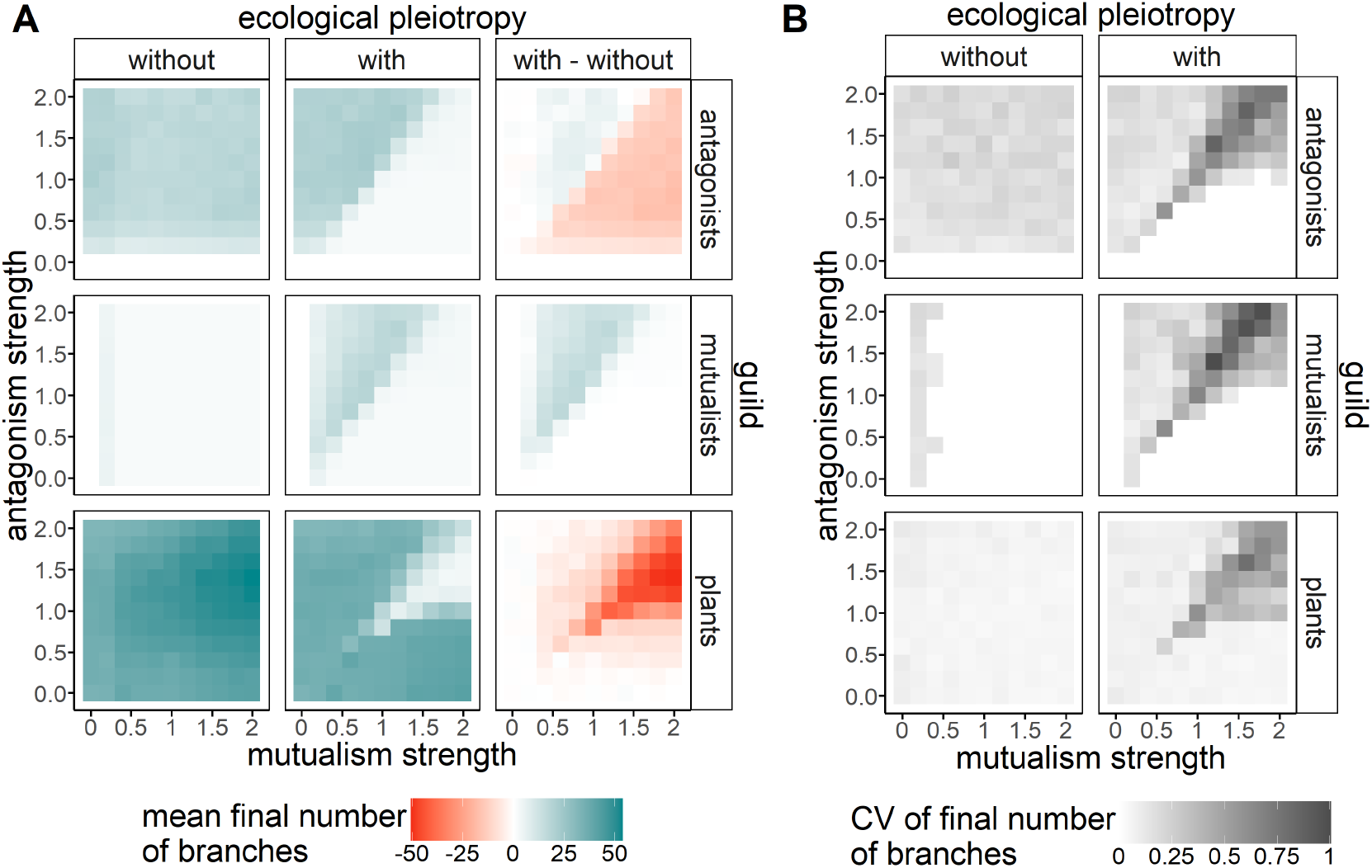
Ecological pleiotropy alters phenotype richness. Mean (A) and coefficient of variation (B) of the number of branches at the end of simulation runs (after 5 billion time steps), for varying strength of mutualism 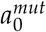 and strength of antagonism 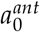, over 20 replicates. Coefficient of variation is the ratio of the standard deviation to the mean. Rows show the different guilds, columns stand for scenario without ecological pleiotropy, with ecological pleiotropy and the difference between the two. All non-varied parameters are chosen according to Table 1.

Without ecological pleiotropy (first column in Figure 4A), the final number of branches is always very low in the mutualist guild, while antagonists and plants reach intermediate to high levels of phenotype richness. This pattern is hardly affected by the strengths of antagonism and mutualism (as long as they are greater than 0, which is necessary for the survival of antagonists/mutualists).

By contrast, with ecological pleiotropy (second column in Figure 4A), the extent of diversification strongly depends on the relative strength of antagonism and mutualism. Confirming the above results from individual simulation runs, all guilds can diversify if antagonism is stronger than mutualism (upper left within each panel). If mutualism is stronger than antagonism, diversification in antagonists and mutualists remains limited (lower right within each panel). Plants still reach a high number of branches in their second trait due to similarity-based competition, explaining the area with high plant phenotype richness in the lower right of the bottom plot. Finally, there is a region where all guilds show a low number of branches, including the plants. This is the case if mutualism and antagonism strength are similar and both large, and the phenomenon is more pronounced when mutualism is slightly stronger than antagonism. For the same parameter combinations, there is high variation among replicates (Fig. 4B), pointing towards recurrent extinction avalanches, as described above. Since the timing of these extinction avalanches is highly stochastic, the number of branches present at a given point in time varies widely between replicates.

The effect of ecological pleiotropy on diversification can only be positive in the case of strong antagonism and only for mutualists and antagonists; otherwise, it is negative or neutral. This effect of ecological pleiotropy is estimated using the difference between the mean final number of branches with and without ecological pleiotropy (third column in Figure 4A). If antagonism is stronger than mutualism (upper left within panels), ecological pleiotropy increases diversification rates in the mutualist guild and, to a lesser extent, also in the antagonistic guild. The final number of plant branches slightly decreases. If mutualism is stronger than antagonism instead (lower right within panels), ecological pleiotropy prevents the diversification in the antagonist guild that would otherwise occur. The number of plant branches slightly decreases, while there is no effect on the diversification of the mutualists. If antagonism and mutualism strength are similar and both high (upper right within panels), ecological pleiotropy creates potential for extinction avalanches, greatly reducing the number of branches in plants and, to a lesser extent, antagonists, compared to the non-pleiotropic case.

The outlined effect of ecological pleiotropy on diversification is mostly robust to changes in various key parameters of the model (see Supplementary Material S2). Increasing the share of similarity-based competition (*α*^*P*^, *α*^*M*^, *α*^*A*^) only intensifies the observed effects (see Fig. S1). The same occurs when the interguild interactions are turned non-obligate for mutualists and antagonists, meaning positive intrinsic net growth rates *r*_*M*_ and *r*_*A*_ (Fig. S2). Changing the relative speed of evolution (by varying the mutation rates *µ*_*P*_, *µ*_*M*_, *µ*_*A*_) can influence the escape-and-chase dynamics between plants and antagonists (and thereby diversification rates), but does not reverse the sign of the effect of ecological pleiotropy either (Fig. S3). Only the variation of mutualism or antagonism kernel width (*σ*_*mut*_, *σ*_*ant*_) can have more intricate, non-monotone consequences for the effect of ecological pleiotropy on diversification (Fig. S4 and Fig. S5). Furthermore, all qualitative results are corroborated by a model version that incorporates an environmental trait optimum, thereby relaxing the assumption of neutrality in the absence of interguild interactions (see Supplementary Material S3).

### Functional diversity is shaped by escape-and-chase dynamics

Finally, we analyse how ecological pleiotropy influences the functional diversity in each of the guilds, measured by the standard deviation of trait values at the end of a simulation run (Fig. 5).

**Figure 5:**
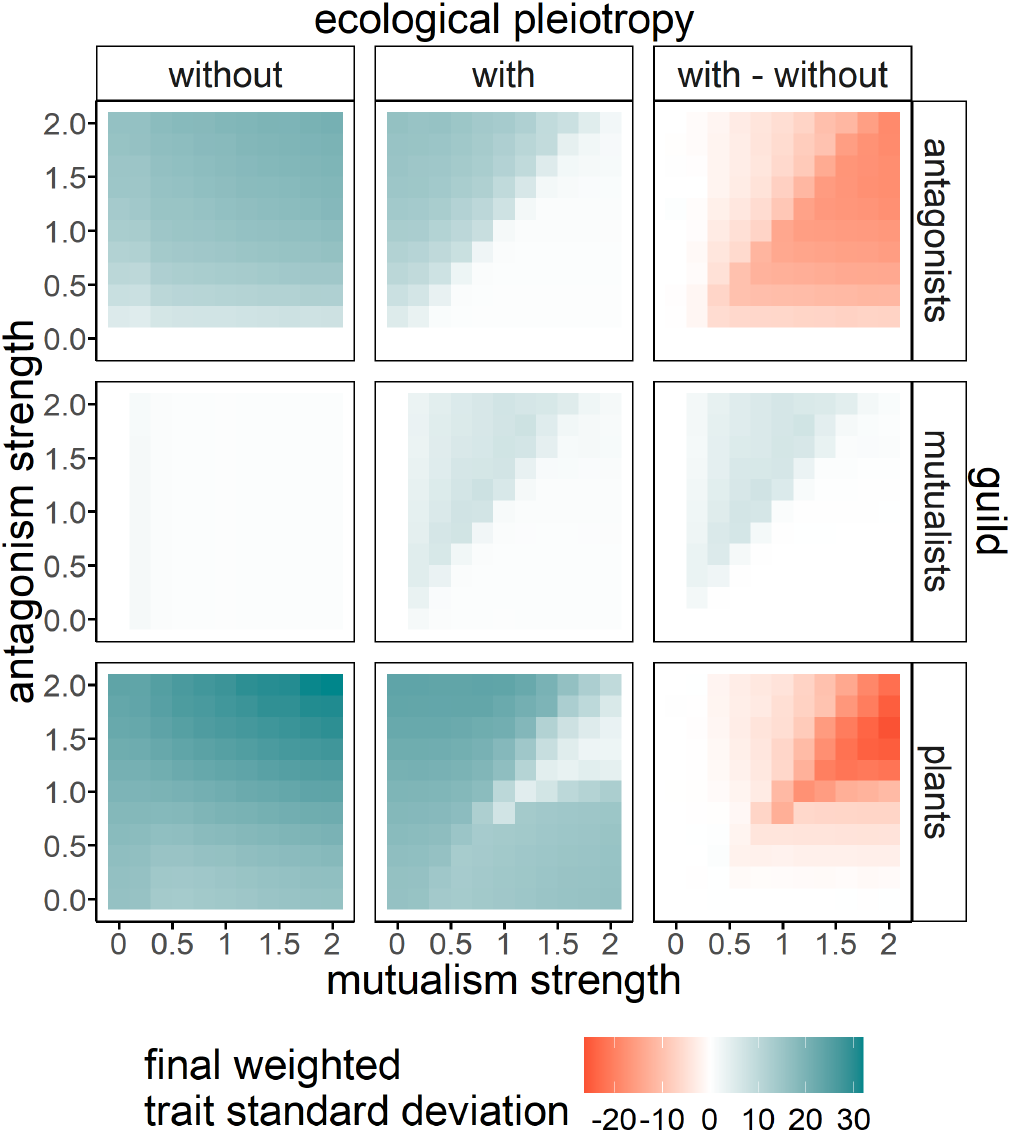
Ecological pleiotropy reduces functional diversity in the antagonistic subnetwork. Mean weighted standard deviation of trait values at the end of simulation runs (after 5 billion time steps), for varying strength of mutualism 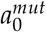 and strength of antagonism 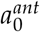, over 20 replicates. Rows show the different guilds, columns stand for scenario without ecological pleiotropy, with ecological pleiotropy and the difference between the two. All non-varied parameters are chosen according to Table 1.

In general, functional diversity patterns align with the patterns in phenotype richness described in the previous subsection. Without ecological pleiotropy, only plants and antagonists diversify. With ecological pleiotropy, in the case of strong mutualism, functional diversity of mutualists and antagonists remains low, but plants may diversify in their second trait (except for the regions where extinction avalanches occur). If antagonism is stronger than mutualism, diversification in all guilds is possible. Plants reach the highest functional diversity, followed by antagonists and mutualists. This reflects the finding from individual simulations that antagonist and mutualist branches stay closer to the centre of the trait space, not following the outermost plant branches, since they are also attracted by the central plant branches.

In contrast to the sheer number of branches, however, functional diversity is additionally shaped by the prevalence of coevolutionary chases of plants by antagonists. Without ecological pleiotropy, escape-and-chase dynamics are common in the antagonistic subnetwork (see Fig. 2D). Their intensity increases with the strength of antagonism, but also with the strength of mutualism since the ecological benefits from mutualism enable higher densities of plants and antagonists, accelerating their evolution. Therefore, functional diversity in plants and antagonists increases with both antagonism and mutualism strength (first column in Fig. 5).

Moreover, different from the results on the number of branches, the effect of ecological pleiotropy on functional diversity (third column in Figure 5) is always negative for antagonists, even for strong antagonism. The reason is that ecological pleiotropy suppresses the escape-and-chase dynamics occurring in the non-pleiotropic case. If mutualism is governed by the same trait as antagonism, runaway selection in plants is counteracted by directional selection towards the mutualists.

## Discussion

In this study, we investigated evolutionary diversification in mutualistic-antagonistic tripartite ecological networks. We found that the presence of ecological pleiotropy significantly alters the evolutionary trajectories and patterns of emerging diversity. Without ecological pleiotropy (i.e., antagonism and mutualism are determined by two uncorrelated plant traits), we observed limited diversification in the mutualistic subnetwork and rapid diversification in the antagonistic subnetwork. Ecological pleiotropy linked the diversification patterns in mutualistic and antagonistic subnetworks. Important implications are the diversification of mutualists in response to antagonism-induced plant diversification, but also the inhibition of antagonist diversification in response to mutualism-induced stabilising selection in plants. Which pattern predominates depends on the relative strength of antagonism and mutualism. Moreover, by suppressing escape-and-chase dynamics, ecological pleiotropy consistently reduces functional diversity in antagonists and plants. Ecological pleiotropy did not in itself generate novel diversification, contradicting the hypothesis that ecological pleiotropy generally enhances plant diversification by creating an ecological trade-off (although we do not rule out the possibility that this mechanism occasionally causes disruptive selection in single plant branches).

Our model corroborates previous results on the effect of single interaction types on evolutionary diversification. Using approaches ranging from quantitative genetics to adaptive dynamics to individual-based modelling and focusing on phenotype richness as well as functional diversity, a variety of theoretical studies have found that mutualism hinders diversification [6, 28, 30, 31]. The reasoning is that mutualism imposes stabilising selection, since both partners benefit from a matching trait. Such an outcome is also consistent with the frequent observation of trait convergence within guilds in mutualistic networks [66, 88, 89, 90]. We acknowledge, however, that various mechanisms not considered in our study (such as pollinator-mediated reproductive isolation) can nevertheless promote diversification in real-world mutualistic interactions [20]. The finding that antagonism fosters diversification is in line with previous work as well. Classical works on plant-insect coevolution suggest that fast divergence of lineages follows a pattern of escape and subsequent radiation [7, 9]. Furthermore, antagonism can directly create disruptive selection in the resource when escape in different directions in the trait space is possible [6, 28, 30]. The antagonist can diversify onto the branches of the resource. Coevolutionary escape-and-chase dynamics further contribute to increased functional diversity by favouring the emergence of extreme phenotypes.

When combining different interaction types, ecological pleiotropy turned out to be an important determinant of the coupling of eco-evolutionary dynamics. Without ecological pleiotropy, one can focus only on one interaction type in isolation to understand diversification patterns. With ecological pleiotropy, one would miss an essential part of the eco-evolutionary story. Based on our results, we could, for instance, speculate that strong herbivory might have caused the unprecedented diversification in angiosperms, which in turn led to the diversification of pollinators onto the available plant resources. Vice versa, the reason for trait convergence in antagonistic guilds could be the presence of another (strong enough) mutualistic interaction mediated by similar plant traits. For example, the phenology of different herbivore species might converge to match plant phenology that has synchronised in response to pollination.

While comprehensive data on diversification in mutualistic-antagonistic systems with and without ecological pleiotropy is not (yet) available, several macroevolutionary studies confirm that the interplay of the two interactions plays a critical role for coevolutionary patterns. For example, Adler et al. [91] use a combination of greenhouse experiments and phylogenetic comparative methods to link increased pollinator reliance with decreased plant defences (nicotine levels) across species of the plant genus *Nicotiana*. By quantifying the correlation of nicotine concentrations in leaf and flower tissues, they can confirm the presence of ecological pleiotropy with respect to pollination and leaf herbivory. In agreement with our results, their findings suggest that strong mutualists can constrain the evolution of antagonism-related traits. Phylogenetic comparative methods have also been applied with an explicit focus on diversification and how it is shaped by biotic interactions [2, 25]; however, to the best of our knowledge, not yet considering multiple types of interaction simultaneously. Another line of research opposes phylogenies of multiple interacting guilds and analyses patterns of co-diversification. For instance, Currie et al. [92] reconstruct the phylogenies of all involved guilds in a tripartite system with ants, their fungal symbionts and fungal parasites. They conclude that the evolution of the mutualism has been shaped by an arms race with the parasite, reflecting the results of our study, assuming a strong enough antagonism. In addition, their study proves that the concepts and questions discussed here enjoy broad applicability beyond plant-related examples. A challenge of their approach is that phylogenetic congruence need not necessarily be the result of coevolution [50, 66].

On the microevolutionary scale, experiments can allow direct investigations of the combined influence of different interactions on selection patterns or evolutionary trajectories. For instance, Gómez [36] finds in a field experiment that the presence of ungulate herbivores disrupted pollinator-mediated selection on the herb *Erysimum mediohispanicum* [see 93, for similar observations in a seed predation/dispersal system]. The author measured correlations between a variety of plant traits and related them to pollinator and herbivore preferences, thereby detecting ecological pleiotropy. Consistent with our results, his empirical study suggests that in the presence of ecological pleiotropy, strong third-party antagonists fundamentally alter mutualistic coevolutionary dynamics. In an evolution experiment, Ramos and Schiestl [38] analyse the change in floral and defence traits over several generations of *Brassica rapa* plants exposed to combinations of herbivory and pollination. They find that flowers do not evolve as much attractiveness to pollinators when herbivores are present, indicating a physiological defence-attractiveness allocation trade-off, which is a form of ecological pleiotropy. Microorganism communities are particularly suitable for experimental evolution approaches due to high densities and short generation times. Harrison et al. [94] use genome sequencing of experimentally evolved bacteria to show that the combination of interactions with phages and plasmids impeded bacterial molecular evolution compared to pairwise scenarios (here, both interactions are negative for the bacteria). Classically employed phytoplankton-zooplankton systems [95] could also be used via the addition of mutualistic bacteria that are increasingly shown to be key in the functioning of phytoplankton communities [96]. While the cited studies do not explicitly focus on diversification, experimental evolution has already been used in this context, for instance, to highlight the effects of interspecific competition on adaptive radiation [97, 98].

Interestingly, several of our simulation runs exhibited occasional extinction avalanches, where a substantial share of the phenotypes in the community went extinct within a short time interval. We hypothesise the extinction avalanches to arise from competitive exclusion, triggered by a mutualist that approaches a plant branch ahead of any antagonist (see Supplementary Material S4 for details). In macroevolutionary studies, periods of rapid extinctions are most often associated with exogenous drivers, such as environmental changes or biological invasions [99, 100]. Our model suggests that, on a local scale, extinction avalanches can also arise from endogenous eco-evolutionary dynamics of biotic interactions only. Similar extinction avalanches have been observed in previous eco-evolutionary models, although the mechanisms driving them presumably vary [52, 101]. In our case, extinction avalanches occurred only with ecological pleiotropy being present, and for similar strength of antagonism and mutualism. Thus, while previous theoretical work suggested that a balance of mutualism and antagonism is essential for ecological stability in mutualistic-antagonistic systems [102, 103, 104], here we show that it creates instability in an evolutionary sense. A thorough investigation to further characterise the phenomenon and its implications will be conducted in a companion study.

For the sake of simplicity and generality, our model naturally makes many assumptions unlikely to be met in nature. In particular, the assumption of trait matching is critical for our results since it shapes the selection patterns generated by antagonism and mutualism. However, empirical studies show that trait matching sometimes only explains a small proportion of variation in interaction partners [67, 68], and, depending on the system under study, other linkage rules might be more appropriate [66, 105, 106]. Changing the linkage rule can fundamentally change the observed diversification patterns, as has been shown in a model by Yoder and Nuismer [28]. Moreover, in nature, the effect of biotic interactions on fitness varies over time, eventually even changing its sign [107, 108]. Such a variation in interaction type would make it impossible to separate the mutualistic and antagonistic guilds, which would have consequences for diversification that are hard to predict. Lastly, biotic interactions are usually not governed by single traits but by a combination of multiple traits. This challenges the idea of ecological pleiotropy being either present or absent, since only some of the traits might affect different interactions simultaneously while others do not, and the degree of phenotypic correlations can vary widely among traits and phenotypes in the system.

In conclusion, our study demonstrates that ecological pleiotropy intertwines the evolutionary trajectories of different interactions, so that they cannot be understood in isolation. In particular, our eco-evolutionary assembly model shows that ecological pleiotropy aligns diversification patterns in mutualistic and antagonistic guilds, although they interact only indirectly via a shared plant guild. Based on our findings, future research can focus on the consequences of ecological pleiotropy for other features of multipartite interaction networks, including their structure and their ecological and evolutionary stability.

## Data and Code Availability

All data and code for this work are available from the Zenodo repository [109]. It can be accessed via the URL https://zenodo.org/records/20613268?token=eyJhbGciOiJIUzUxMiJ9.eyJpZCI6IjgxNGEwMmMyLWNiOWEtNDZjZS04MDIxLTAyZmNjMmRlZDA5ZCIsImRhdGEiOnt9LCJyYW5kb20iOiJmNzZlZTBiYmExZTY2NmY0MjFiYmIyMzBlMjkyYTdhMyJ9.aSk0Ppb68L2nPvU8dd4uK0T8MFGSwBNW--KlL_0ck8dCf7WTbxyO_ev8VE2T2u4MR6RPX-tWQkWd0cU5RNiT_Q.

