## Supplementary Material for "Between Friends and Foes: Evolutionary Diversification in Mutualistic-Antagonistic Networks"

Felix Jäger[1,2,*]

Nicolas Loeuille[3]

Youssef Yacine[4]

Korinna T. Allhoff[1,2,5,6]

1. Department of Eco-Evolutionary Modelling, University of Hohenheim, Stuttgart, Germany;

2. Computational Science Hub (CSH), University of Hohenheim, Stuttgart, Germany;

3. Sorbonne Université, Université Paris Cité, Univ Paris Est Créteil, CNRS, IRD, INRAE, Institut d'Écologie et des Sciences de l'Environnement de Paris (iEES-Paris), Paris, France;

4. MARBEC, Université de Montpellier, CNRS, Ifremer, IRD, Montpellier, France;

5. KomBioTa - Center for Biodiversity and Integrative Taxonomy, University of Hohenheim & State Museum of Natural History, Stuttgart, Germany;

6. Cluster of Excellence GreenRobust, University of Hohenheim, Stuttgart, Germany;

#### S1 Similarity-based competition is necessary for diversification

In this section, we use the framework of Adaptive Dynamics to analytically show that some extent of similarity-based competition is necessary for evolutionary diversification to occur in our model. The underlying intuition is that with only trait-independent competition, the difference between interspecific and intraspecific competition would be too small to allow the coexistence of the resident type and the mutant type after a mutation [1]. This result is corroborated by the numerical simulations in Supplementary Material S2 which do not exhibit any diversification without similarity-based competition.

We consider the initial system with only one phenotype in each guild. The only possible evolutionary singularity occurs with perfect matching of traits in both the antagonistic and the mutualistic interaction, otherwise the antagonist or the mutualist would experience directional selection towards the plant. It is clear that in such a situation only the plant phenotype potentially experiences disruptive selection, as mutualist and antagonist benefit from the trait matching. Similarly, it is clear that the mutualism does not contribute to the disruptive selection in the plant, since the plant benefits from the mutualistic trait matching. Therefore, it suffices to look at the reduced system of only plant and antagonist and focus only on potential branching in the plant. We can further assume that the plant has only one trait $t^P$, since the second plant trait is not under selection without similarity-based competition.

Under these assumptions, the equations describing plant and antagonist population dynamics are

$$\frac{dP}{dt} = P\left(r^P - c^P(t^P, t^P)P - \frac{a^{ant}(t^P, t^A)A}{1 + h_A a^{ant}(t^P, t^A)P}\right) \tag{S1}$$

$$\frac{dA}{dt} = A\left(r^A - c^A(t^A, t^A)A + \frac{e_A a^{ant}(t^P, t^A)P}{1 + h_A a^{ant}(t^P, t^A)P}\right), \tag{S2}$$

where

$$c^P(t_i^P, t_j^P) = c_0^P \left( (1 - \alpha^P) + \alpha^P \exp\left( \frac{-(t_i^P - t_j^P)^2}{2\sigma_{PC}^2} \right) \right) \tag{S3}$$

$$c^A(t_i^A, t_j^A) = c_0^A \left( (1 - \alpha^A) + \alpha^A \exp\left( \frac{-(t_i^A - t_j^A)^2}{2\sigma_{AC}^2} \right) \right) \tag{S4}$$

$$a^{ant}(t_i^P, t_j^A) = a_0^{ant} \exp\left( \frac{-(t_i^P - t_j^A)^2}{2\sigma_{ant}^2} \right) \tag{S5}$$

Following the framework of Adaptive Dynamics, we assume the plant and antagonist densities to be at a (feasible) equilibrium $P^*$ and $A^*$. We quantify the invasion fitness of a plant mutant $P_m$ with a slightly different trait value $t_m^P$ than the resident trait value $t_r^P$, that is, the per-capita growth rate of the mutant when rare:

$$f(t_r^P, t_m^P) = r^P - c^P(t_m^P, t_r^P)P^* - \frac{a^{ant}(t_m^P, t^A)A^*}{1 + h_A a^{ant}(t_r^P, t^A)P^*} \tag{S6}$$

$$= r^P - c^P(t_m^P, t_r^P)P^* - \frac{a^{ant}(t_m^P, t^A)}{a^{ant}(t_r^P, t^A)} \cdot \left( r^P - c_0^P P^* \right). \tag{S7}$$

Here we have used that $A^*$ and $P^*$ render the right hand side of equation S1 zero. Note that $A^*$ and $P^*$ depend on $t_r^P$ and $t^A$, but not on $t_m^P$.

The selection gradient for the plant is

$$\left. \frac{\partial f(t_r^P, t_m^P)}{\partial t_m^P} \right|_{t_m^P = t_r^P} = \frac{t_r^P - t^A}{\sigma_{ant}^2} \cdot \left( r^P - c_0^P P^* \right), \tag{S8}$$

using that $\frac{\partial c^P(t_m^P, t_r^P)}{\partial t_m^P}\big|_{t_m^P = t_r^P} = 0$ and $\frac{\partial a^{ant}(t_m^P, t^A)}{\partial t_m^P}\big|_{t_m^P = t_r^P} = -\frac{t_r^P - t^A}{\sigma_{ant}^2} \cdot a^{ant}(t_r^P, t^A)$. The unique evolutionarily singular strategy, where the selection gradient vanishes, is $t_r^P = t^A$. We want to check if evolutionary branching can arise from this singularity. One necessary condition is the possible coexistence of the two branches right after the branching (often called protected dimorphism). In the framework of Adaptive Dynamics, the criterion we need to check is the following mutual invasibility criterion [2]:

$$\left(\frac{\partial^2 f(t_r^P, t_m^P)}{\partial (t_m^P)^2} + \frac{\partial^2 f(t_r^P, t_m^P)}{\partial (t_r^P)^2}\right)\Bigg|_{t_m^P = t_r^P = t^A} > 0 \tag{S9}$$

Using that

$$\frac{\partial^2 c(t_m^P, t_r^P)}{\partial (t_m^P)^2}\Bigg|_{t_m^P = t_r^P} = -\frac{c_0^P \alpha^P}{\sigma_{PC}^2} \tag{S10}$$

and

$$\frac{\partial^2 a(t_m^P, t^A)}{\partial (t_m^P)^2}\Bigg|_{t_m^P = t^A} = -\frac{a_0^{ant}}{\sigma_{ant}^2}, \tag{S11}$$

we get

$$\frac{\partial^2 f(t_r^P, t_m^P)}{\partial (t_m^P)^2}\Bigg|_{t_m^P = t_r^P = t^A} = \frac{c_0^P \alpha^P}{\sigma_{PC}^2} \cdot P^* + \frac{r^P - c_0^P P^*}{\sigma_{ant}^2}. \tag{S12}$$

Note that this curvature is positive (since $r^P - c_0^P P^* = \frac{a_0^{ant} A^*}{1 + h_A a_0^{ant} P^*}$), meaning that the plant is at a fitness minimum, even without similarity-based competition. However, we further have

$$\frac{\partial^2 f(t_r^P, t_m^P)}{\partial (t_r^P)^2} = \frac{c_0^P \alpha^P}{\sigma_{PC}^2} \cdot P^* - \frac{r^P - c_0^P P^*}{\sigma_{ant}^2}, \tag{S13}$$

which yields

$$\left(\frac{\partial^2 f(t_r^P, t_m^P)}{\partial (t_m^P)^2} + \frac{\partial^2 f(t_r^P, t_m^P)}{\partial (t_r^P)^2}\right)\Bigg|_{t_m^P = t_r^P = t^A} = \frac{2 c_0^P \alpha^P}{\sigma_{PC}^2} \cdot P^*. \tag{S14}$$

This is only positive, if $\alpha^P$ is positive. This means some extent of similarity-based competition in the plants is necessary for the first branching event in our model. Further simulations (not shown) prove that similarity-based competition is also required in the mutualists to potentially allow subsequent diversification of the mutualists. While this is not the case for antagonists, we still assume similarity-based competition in all guilds for comparability.

### S2 Robustness analysis

In this section, we outline how varying several key parameters of the model (namely the proportions of similarity-based competition $\alpha_P$, $\alpha_M$ and $\alpha_A$, the net intrinsic growth rates of mutualists and antagonists $r_M$ and $r_A$, the ratio of the mutation rates $\mu_P$, $\mu_M$ and $\mu_M$, and the mutualism/antagonism kernel widths $\sigma_{mut}$ and $\sigma_{ant}$) influences the effect of ecological pleiotropy on diversification. We focus on the final number of branches as a measure of diversity, while the standard deviation of diversity between replicates (especially in plants) serves as an indicator for the presence of extinction avalanches. We compare simulations with and without ecological pleiotropy. To cover the different possible qualitative outcomes described in the main text, we compare scenarios with stronger antagonism, equal strength of antagonism and mutualism, and stronger mutualism.

Diversification in all guilds increases with increasing proportion of similarity-based competition, intensifying the effects of ecological pleiotropy (see Figure S1). Without similarity-based competition ($\alpha^P = \alpha^M = \alpha^A = 0$), we observe no diversification in any of the guilds, both with and without ecological pleiotropy, corroborating the analytical result from Supplementary Material S1. With intermediate values of $\alpha^P, \alpha^M, \alpha^A$ (starting from 0.2), diversification in the antagonistic subnetwork and, in the presence of ecological pleiotropy and strong enough antagonism, also in the mutualistic subnetwork becomes possible. The sign of the effect of ecological pleiotropy (third column of the figure) does not change in any of the guilds for varying $\alpha^P = \alpha^M = \alpha^A$ (only the effect for antagonists in case of strong antagonism can get slightly negative). This means that the patterns described in the main text are qualitatively robust to varying strength of similarity-based competition. With high values of $\alpha$ (starting from 0.5), simulations tend to stop prematurely since the maximum number of phenotypes is reached in one of the guilds (missing values in the figure). With an increased share of similarity-based competition, the standard deviation of the number of plant branches across replicates decreases in the case of ecological pleiotropy and equal strength of mutualism and antagonism. This indicates that extinction avalanches cease to exist if similarity-based competition is strengthened relative to trait-independent competition.

The general patterns observed in the main text can also be found for non-obligate interactions (Figure S2). If the intrinsic net growth rates $r^M$ and $r^A$ of mutualists and antagonists are positive, meaning that the populations can also grow in the absence of the focal plant resources, diversification rates of mutualists and antagonists are increased in general via increased densities. This intensifies the effect of ecological pleiotropy, while leaving its sign unchanged. If, however, $r^M$ and $r^A$ are decreased below $-0.1$, diversification in antagonists and mutualists remains limited both with and without ecological pleiotropy (due to reduced densities), and the effect of ecological pleiotropy no longer unfolds.

Varying the relative mutation rate of plants, antagonists and mutualists mostly does not alter the effect of ecological pleiotropy on diversification (Figure S3). A qualitative deviation of the main patterns is only observed for increased plant mutation rate. In that case, plants tend to escape antagonists and mutualists, leaving them at low densities and low diversification rates, regardless of the presence of ecological pleiotropy. Hence, ecological pleiotropy has almost no effect on mutualist and antagonist diversification. However, it can reduce plant diversification via extinction avalanches (indicated by large error bars) in the case of equal strengths of mutualism and antagonism, or, to a lesser extent, in the case of strong antagonism. By contrast, extinction avalanches are prevented by fast antagonist evolution, indicated by low variation in plant diversity between replicates, even with ecological pleiotropy and equal strength of mutualism and antagonism. Accordingly, the effect of pleiotropy on plant diversification is not as negative as in the baseline scenario. Instead, the negative effect of ecological pleiotropy on antagonist diversity is more pronounced than in the baseline scenario (if mutualism is at least as strong as antagonism). This is due to the fact that without ecological pleiotropy, antagonists can reach very high levels of diversity due to increased speed of evolution. Increasing the mutation rate of mutualists has almost no consequences for diversity in any of the scenarios.

The sign of the effect of ecological pleiotropy remains largely unchanged also under varying mutualism kernel width $\sigma_{mut}$ (Figure S4). Increasing mutualism kernel width increases the strength of mutualistic interactions (see equation 4 in the main text), hence the densities of plants and mutualists. Therefore, it tends to increase diversification in plants and mutualists both with and without ecological pleiotropy. If the mutualism niche is too narrow ($\sigma_{mut} < 2.5$), mutualists do not diversify at all, even with ecological pleiotropy and strong antagonism. This eliminates the potential positive effect of ecological pleiotropy for mutualist diversification and can be attributed to stronger stabilising selection. With very narrow mutualism kernel widths (0.5), stabilising selection on plants and mutualists is so strong that even the antagonists cannot diversify in case of ecological pleiotropy and strong antagonism, translating into a negative effect of ecological pleiotropy for antagonist diversification. Interestingly, significant variation in plant diversity between replicates and a strong negative effect of ecological pleiotropy for plant diversity are only found for intermediate $\sigma_{mut}$. This indicates that extinction avalanches are prevented both by increasing and decreasing mutualism kernel width.

The potential positive effect of ecological pleiotropy on mutualist diversification, as observed in the main analysis, occurs only for intermediate antagonism kernel width $\sigma_{ant}$ (Figure S5). If $\sigma_{ant} \leq 1$ or $\sigma_{ant} \geq 3$, diversification in the mutualistic guild remains limited even in the case of ecological pleiotropy and strong antagonism. Moreover, extinction avalanches are prevented in this case, also with equal strengths of antagonism and mutualism. Other than that, changing antagonism kernel width does not affect the main patterns qualitatively. Increasing $\sigma_{ant}$ in general increases diversification rates of antagonists and plants due to stronger antagonistic interactions (see equation 5 in the main text), hence intensifies the potential negative effect of ecological pleiotropy on antagonist diversification.

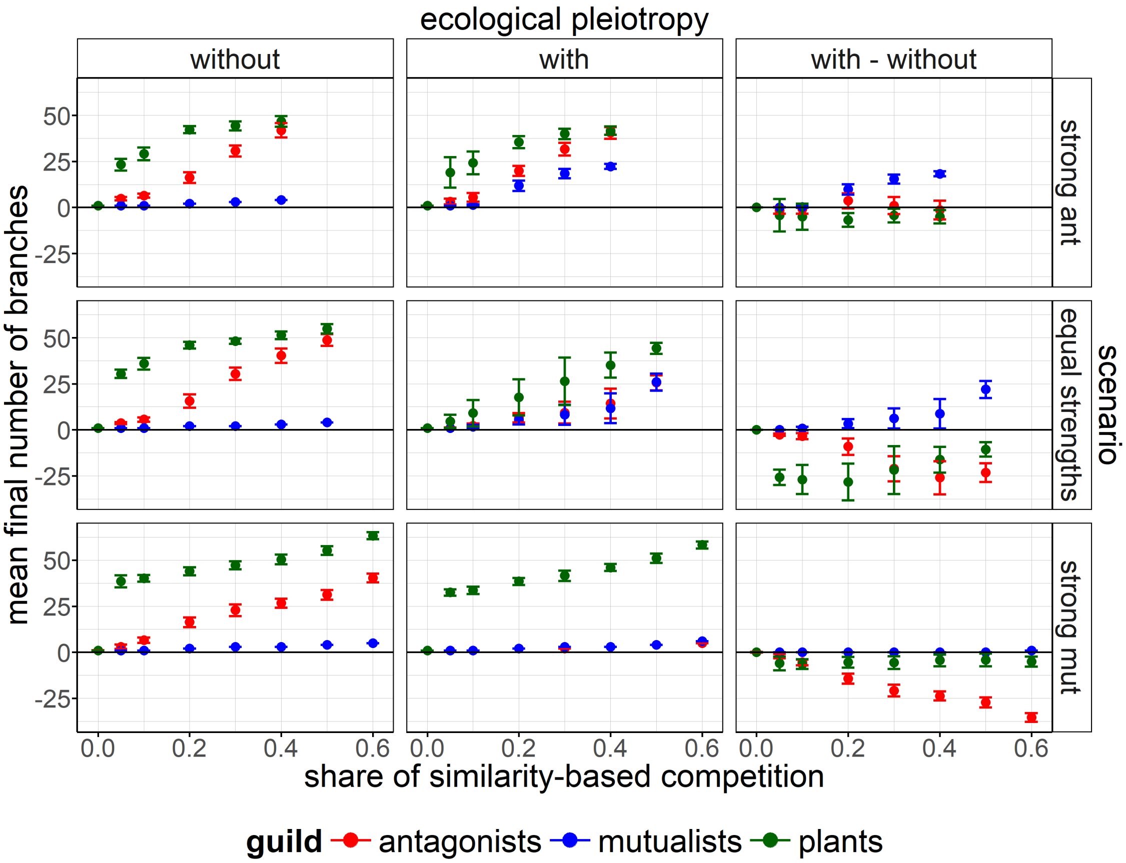


Figure S1: Final number of branches for varying proportion of similarity-based competition $\alpha^P = \alpha^M = \alpha^A$. Rows show 3 different scenarios: (i) strong antagonism ($a_0^{mut} = 0.5$, $a_0^{ant} = 1.5$), (ii) equal strengths ($a_0^{mut} = a_0^{ant} = 1$) and (iii) strong mutualism ($a_0^{mut} = 1.5$, $a_0^{ant} = 0.5$). Columns stand for scenario without ecological pleiotropy, with ecological pleiotropy and the difference between the two. A mean is taken over 20 replicates, error bars show the respective standard deviation. All non-varied parameters are chosen according to Table 1 from the main text (the baseline scenario corresponds to $\alpha^P = \alpha^M = \alpha^A = 0.2$). Missing values correspond to simulation runs that ended prematurely because the maximum number of phenotypes was reached in one of the guilds.

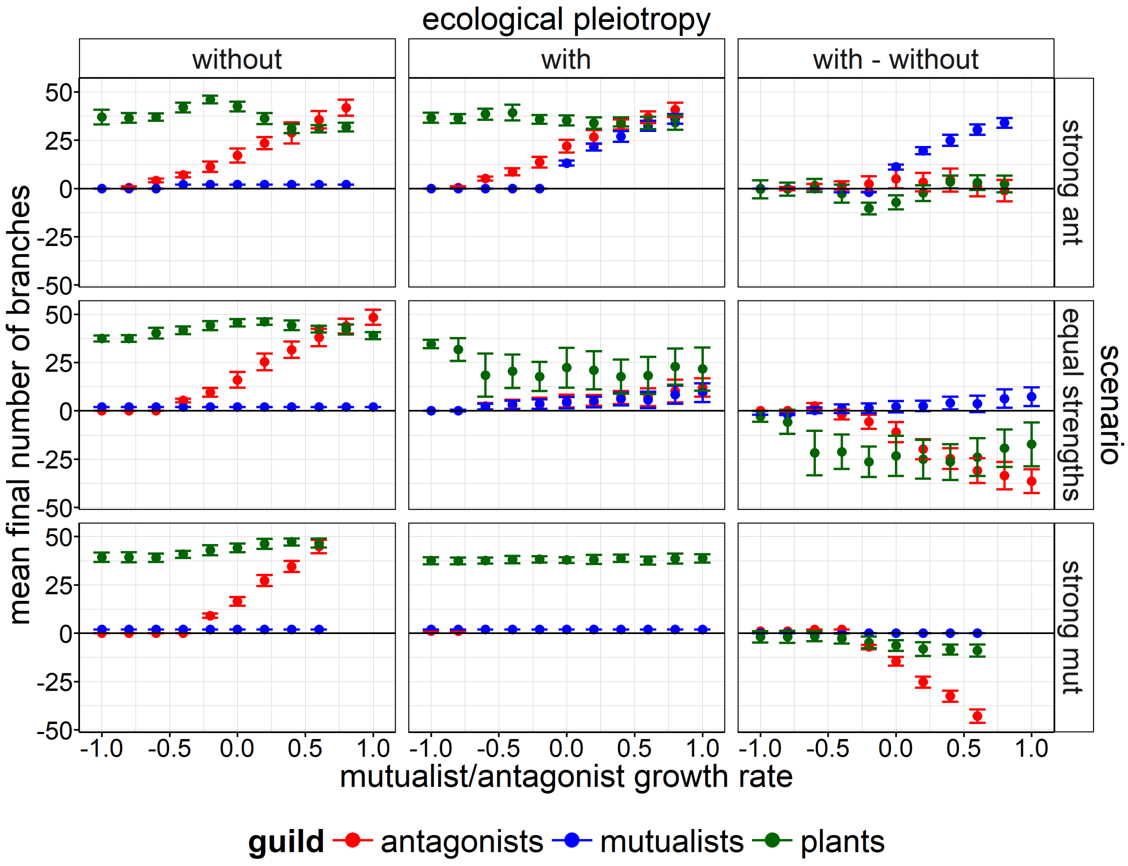


Figure S2: Final number of branches for varying antagonist and mutualist intrinsic net growth rate $r^A = r^M$. Rows show 3 different scenarios: (i) strong antagonism ($a_0^{mut} = 0.5$, $a_0^{ant} = 1.5$), (ii) equal strengths ($a_0^{mut} = a_0^{ant} = 1$) and (iii) strong mutualism ($a_0^{mut} = 1.5$, $a_0^{ant} = 0.5$). Columns stand for scenario without ecological pleiotropy, with ecological pleiotropy and the difference between the two. Mean is taken over 20 replicates, error bars show the respective standard deviation. All non-varied parameters are chosen according to Table 1 from the main text (the baseline scenario corresponds to $r^M = r^A = -0.01$). Missing values correspond to simulation runs that ended prematurely because the maximum number of phenotypes was reached in one of the guilds.

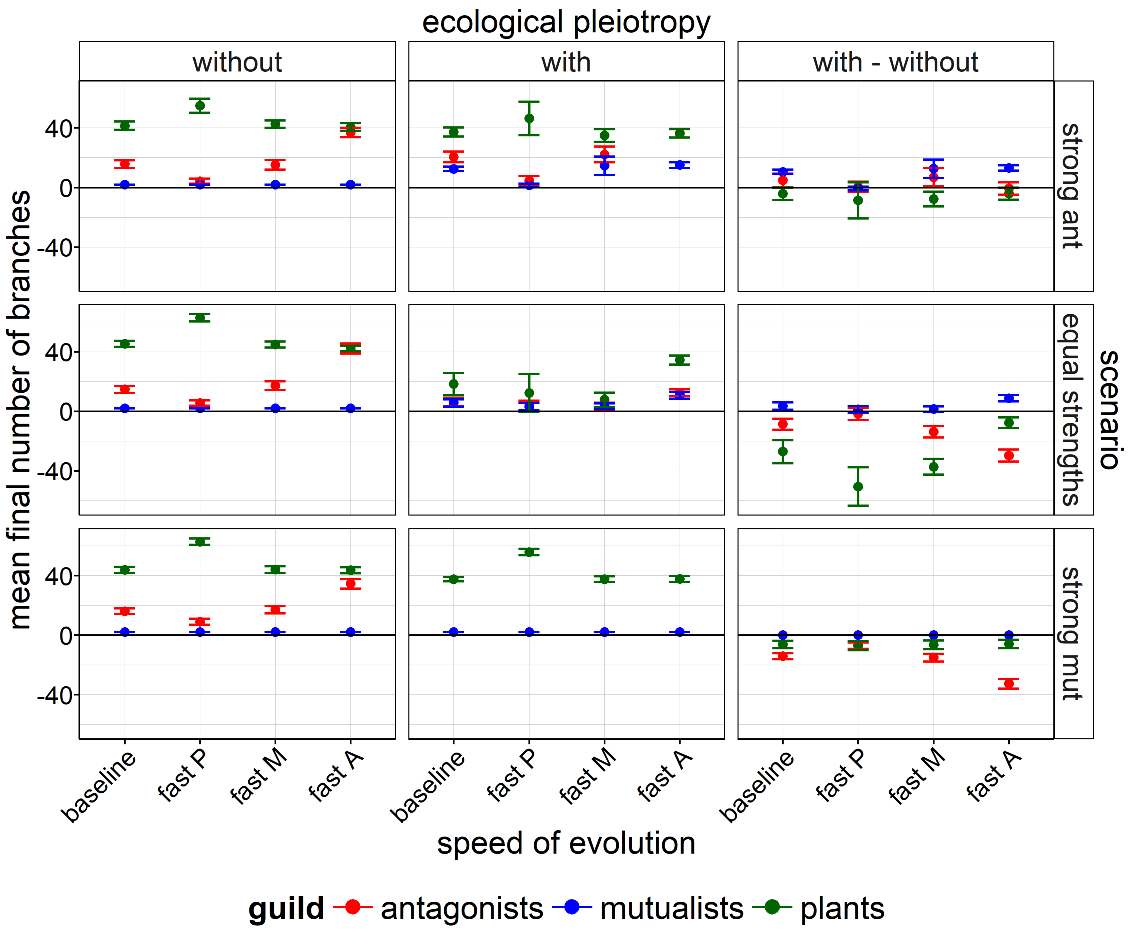


Figure S3: Final number of branches for varying relative speed of evolution. Simulations are run for the baseline parametrisation and with either $\mu_P = 4 \cdot 10^{-6}$ ('fast P'), $\mu_M = 4 \cdot 10^{-6}$ ('fast M'), or $\mu_H = 4 \cdot 10^{-6}$ ('fast H'). Rows show 3 different scenarios: (i) strong antagonism ($a_0^{mut} = 0.5$, $a_0^{ant} = 1.5$), (ii) equal strengths ($a_0^{mut} = a_0^{ant} = 1$) and (iii) strong mutualism ($a_0^{mut} = 1.5$, $a_0^{ant} = 0.5$). Columns stand for scenario without ecological pleiotropy, with ecological pleiotropy and the difference between the two. Mean is taken over 20 replicates, error bars show the respective standard deviation. All non-varied parameters are chosen according to Table 1 from the main text.

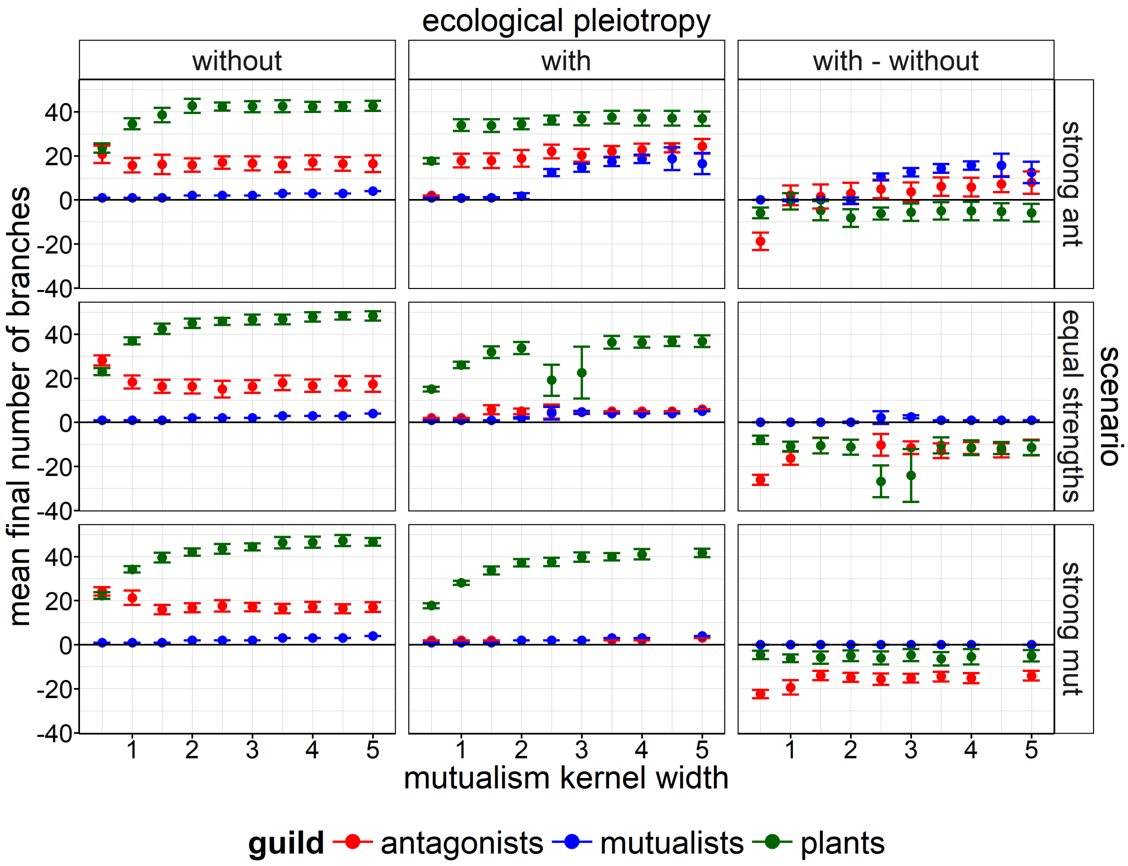


Figure S4: Final number of branches for varying mutualism kernel width $\sigma_{mut}$. Rows show 3 different scenarios: (i) strong antagonism ($a_0^{mut} = 0.5$, $a_0^{ant} = 1.5$), (ii) equal strengths ($a_0^{mut} = a_0^{ant} = 1$) and (iii) strong mutualism ($a_0^{mut} = 1.5$, $a_0^{ant} = 0.5$). Columns stand for scenario without ecological pleiotropy, with ecological pleiotropy and the difference between the two. Mean is taken over 20 replicates, error bars show the respective standard deviation. All non-varied parameters are chosen according to Table 1 from the main text (the baseline scenario corresponds to $\sigma_{mut} = 2.5$). Missing values correspond to simulation runs that ended prematurely because the maximum number of phenotypes was reached in one of the guilds.

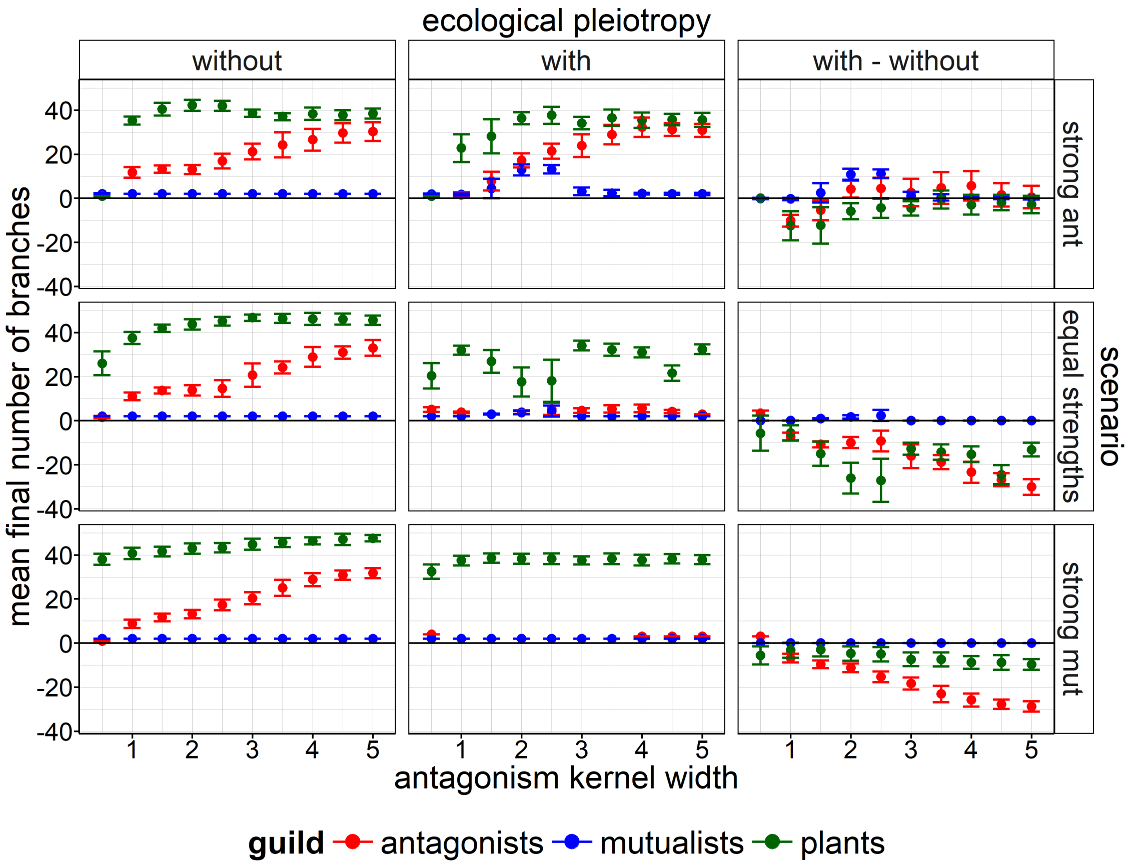


Figure S5: Final number of branches for varying antagonism kernel width $\sigma_{ant}$. Rows show 3 different scenarios: (i) strong antagonism ($a_0^{mut} = 0.5$, $a_0^{ant} = 1.5$), (ii) equal strengths ($a_0^{mut} = a_0^{ant} = 1$) and (iii) strong mutualism ($a_0^{mut} = 1.5$, $a_0^{ant} = 0.5$). Columns stand for scenario without ecological pleiotropy, with ecological pleiotropy and the difference between the two. Mean is taken over 20 replicates, error bars show the respective standard deviation. All non-varied parameters are chosen according to Table 1 from the main text (the baseline scenario corresponds to $\sigma_{ant} = 2.5$). Missing values correspond to simulation runs that ended prematurely because the maximum number of phenotypes was reached in one of the guilds.

### S3 Model version with environmental trait optimum

The model, as presented in the main text, assumes full neutrality in the absence of ecological interactions, meaning that no trait value has a selective advantage over others. In reality, trait values are always subject to additional environmental constraints. For instance, flower size cannot be arbitrarily high but is limited by physiological constraints independent of the interaction with a pollinator. In this section, to ensure robustness of our main results, we consider an extended version of the model by adding such environmental constraints. In particular, we let the plant intrinsic net growth rate $r^P$ depend on the respective plant traits. We assume $r^P$ to follow a quadratic function which attains its maximum when both trait values are 0. Phenotypes that deviate too much from this optimal trait value suffer from reduced intrinsic growth.

$$r^P(t_1^P, t_2^P) = r_0^P - b_{env} \cdot \Big((t_1^P)^2 + (t_2^P)^2\Big) \tag{S15}$$

Here, $r_0^P$ is the maximal intrinsic plant growth rate and $b_{env}$ modulates how fast the net intrinsic growth rate declines when traits move away from the optimum. Note that we do not introduce a similar constraint for mutualists and antagonists, but their trait values are implicitly also constrained since they cannot survive without the interaction with the plants ($r^M$ and $r^A$ are negative).

Simulations with this modified model version corroborate the results from the main text on the influence of ecological pleiotropy on diversification and the dependency on mutualism and antagonism strength (cf. Figure S6). The main difference is that, in cases of high diversification, the trait space does not keep getting filled up with branches indefinitely as was the case in the original model version. Instead, branches are constrained to a region centred around the optimal trait value 0 where the plant intrinsic net growth rate is large enough for survival. Consequently, the number of branches saturates in all guilds.

When the system approaches such saturation, fitness differences between mutant and resident phenotypes eventually become very small and it may take a long time until phenotypes get outcompeted. This weak selection results in an accumulation of phenotypes, especially in the antagonist guild, which explains why many simulation runs in Figure S6 stop prematurely since the maximum number of phenotypes is exceeded.

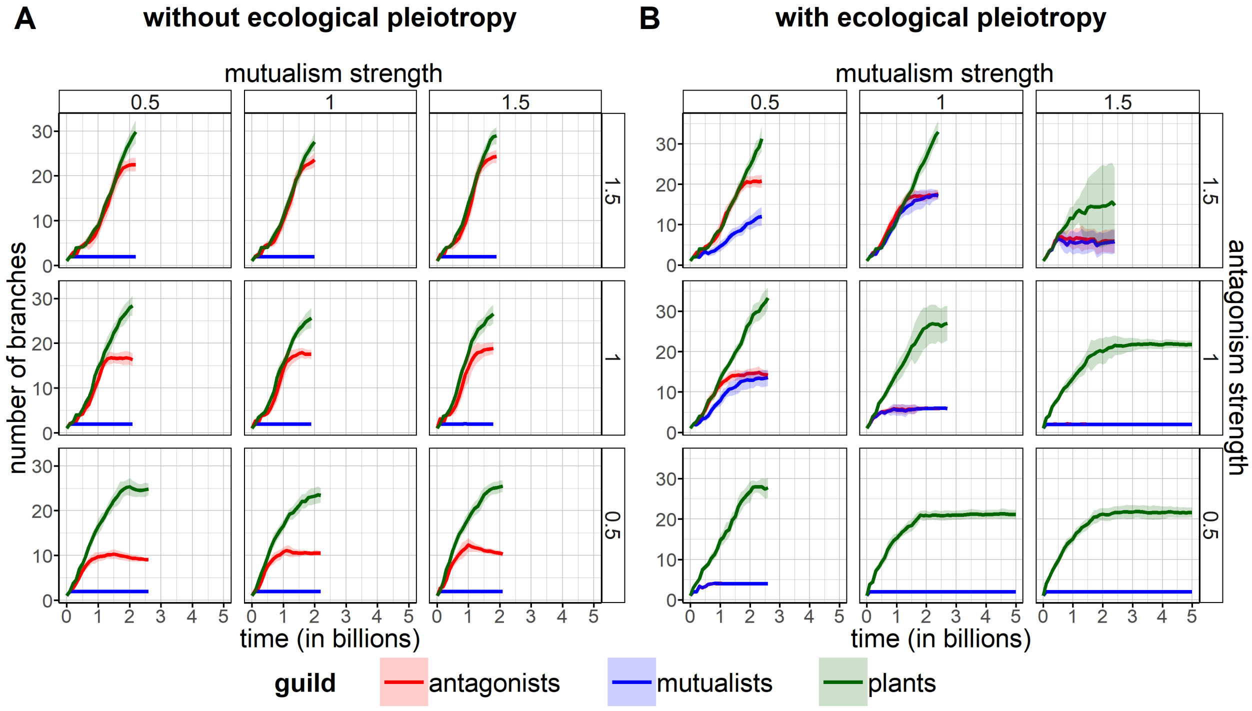


Figure S6: **Diversification in a model version with environmental trait optimum.** Time series show the mean number of branches for varying strength of mutualism $a_0^{mut}$ and strength of antagonism $a_0^{ant}$ over 20 replicates, without ecological pleiotropy (A) and with ecological pleiotropy (B). Shaded areas visualise the standard deviation. Simulations were run with $r_0^P = 10$ and $b_{env} = 0.002$, all other parameters were chosen according to Table 1 from the main text. Initial trait values were set to 0, maximising initial plant net intrinsic growth rate. Missing values correspond to situations where at least one replicate run stopped because the maximum number of phenotypes was reached in one of the guilds.

### S4 Mechanism of extinction avalanches

We hypothesise the extinction avalanches observed in our study to arise from competitive exclusion, triggered by outer mutualist and antagonist phenotypes that approach an outer plant in a specific order. We illustrate the underlying mechanism based on an example simulation run shown in Fig. S7. Starting with a diverse network, one of the outermost mutualist branches approaches an outer plant branch ahead of the closest antagonist (in the figure, the lowest mutualist distances itself slightly from the lowest antagonist after around 1.59 billion time steps). Subsequently, the mutualist and its plant interaction partner benefit from a strong mutualistic interaction that is relatively undisturbed by antagonists and can achieve higher densities. Eventually, the focal mutualist can outcompete the other mutualists within a short time period (at around 1.6 billion time steps in the figure). Similarly, the closest antagonist benefits from the high plant density and can outcompete the other antagonists. Now the remaining antagonist and mutualist branches engage in a race where the mutualist is attracted by available plant branches further away from the antagonist, while the antagonist is attracted by the same plants as soon as the mutualist approaches them and their densities rise. During this process, the plant branch closest to the remaining mutualist has the potential to outcompete other plant branches if the antagonist is far enough away (after approx. 1.66 billion time steps in the example). However, this subsequent extinction avalanche of plant branches does not necessarily occur; e.g., in the bottom left panel of Figure **??** only mutualists and antagonists go extinct.

This hypothesis is in line with the result that extinction avalanches only happen with ecological pleiotropy and with a balance of antagonism and mutualism strength. If mutualism is too strong (or ecological pleiotropy absent), diversification in the mutualistic subnetwork remains limited in the first place, preventing the possibility of extinction avalanches. If antagonism is too strong, instead, high densities of antagonists speed up antagonist evolution since mutation probability depends on density in our model (see Methods). Therefore, the outer plant branches are always first reached by antagonists instead of mutualists in the coevolutionary process, preventing an extinction avalanche according to the described pattern. Similarly, they are prevented by increased antagonist mutation rates (see robustness analysis). Our hypothesis is further supported by the fact that extinction avalanches cease to occur when competition is purely similarity-based (see Figure S1), ruling out the possibility of competitive exclusion over long distances in the trait space.

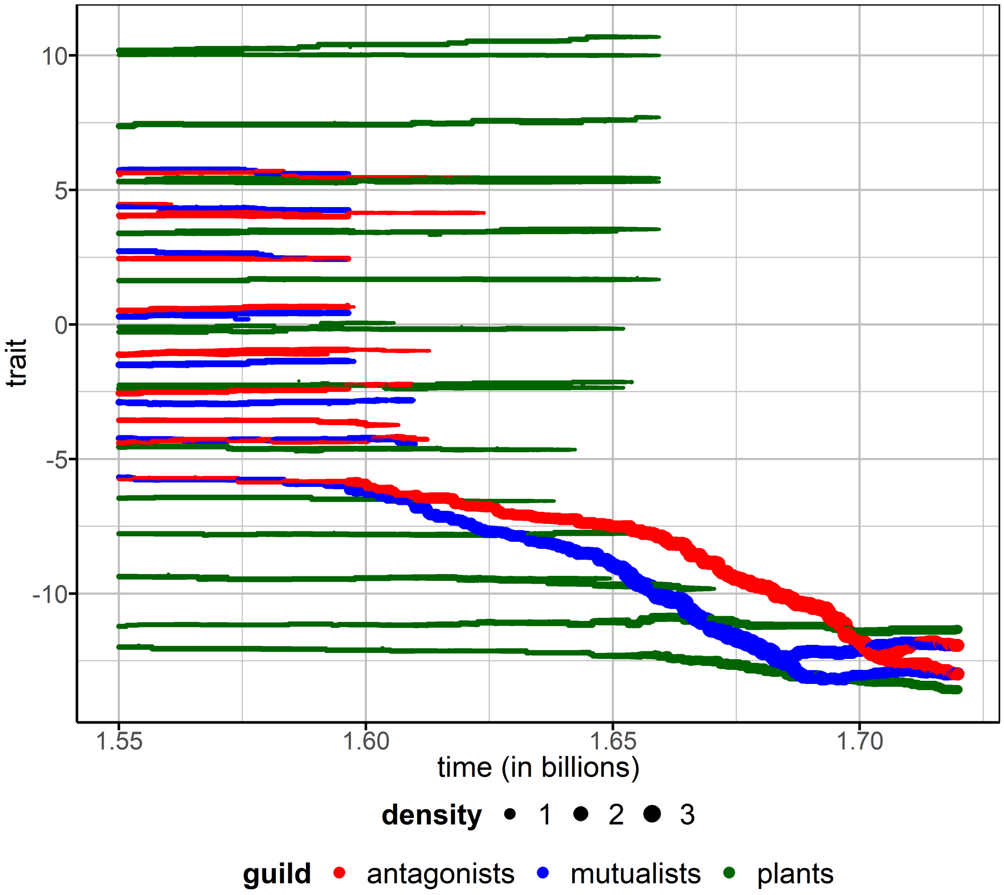


Figure S7: Time series showing the antagonist, mutualist and first plant trait in a simulation run with ecological pleiotropy. Line width represents density of the phenotypes. All parameters are chosen according to Table 1 from the main text. To focus on the extinction avalanche, only the period between 1.55 and 1.72 billion time steps is shown.
